# A Comparison of Methods to Harmonize Cortical Thickness Measurements Across Scanners and Sites

**DOI:** 10.1101/2021.09.22.461242

**Authors:** Delin Sun, Gopalkumar Rakesh, Courtney C. Haswell, Mark Logue, C. Lexi Baird, Brian M. O’Leary, Andrew S. Cotton, Hong Xie, Marijo Tamburrino, Tian Chen, Emily L. Dennis, Neda Jahanshad, Lauren E. Salminen, Sophia I. Thomopoulos, Faisal Rashid, Christopher R. K. Ching, Saskia B. J. Koch, Jessie L. Frijling, Laura Nawijn, Mirjam van Zuiden, Xi Zhu, Benjamin Suarez-Jimenez, Anika Sierk, Henrik Walter, Antje Manthey, Jennifer S. Stevens, Negar Fani, Sanne J.H. van Rooij, Murray Stein, Jessica Bomyea, Inga K. Koerte, Kyle Choi, Steven J.A. van der Werff, Robert R. J. M. Vermeiren, Julia Herzog, Lauren A. M. Lebois, Justin T. Baker, Elizabeth A. Olson, Thomas Straube, Mayuresh S. Korgaonkar, Elpiniki Andrew, Ye Zhu, Gen Li, Jonathan Ipser, Anna R. Hudson, Matthew Peverill, Kelly Sambrook, Evan Gordon, Lee Baugh, Gina Forster, Raluca M. Simons, Jeffrey S. Simons, Vincent Magnotta, Adi Maron-Katz, Stefan du Plessis, Seth G. Disner, Nicholas Davenport, Daniel W. Grupe, Jack B. Nitschke, Terri A. deRoon-Cassini, Jacklynn M. Fitzgerald, John H. Krystal, Ifat Levy, Miranda Olff, Dick J. Veltman, Li Wang, Yuval Neria, Michael D. De Bellis, Tanja Jovanovic, Judith K. Daniels, Martha Shenton, Nic J.A. van de Wee, Christian Schmahl, Milissa L. Kaufman, Isabelle M. Rosso, Scott R. Sponheim, David Bernd Hofmann, Richard A. Bryant, Kelene A. Fercho, Dan J. Stein, Sven C. Mueller, Bobak Hosseini, K. Luan Phan, Katie A. McLaughlin, Richard J. Davidson, Christine L. Larson, Geoffrey May, Steven M. Nelson, Chadi G. Abdallah, Hassaan Gomaa, Amit Etkin, Soraya Seedat, Ilan Harpaz-Rotem, Israel Liberzon, Theo G.M. van Erp, Xin Wang, Paul M. Thompson, Rajendra A. Morey

## Abstract

Results of neuroimaging datasets aggregated from multiple sites may be biased by site-specific profiles in participants’ demographic and clinical characteristics, as well as MRI acquisition protocols and scanning platforms. We compared the impact of four different harmonization methods on results obtained from analyses of cortical thickness data: (1) linear mixed-effects model (LME) that models site-specific random intercepts (LME_INT_), (2) LME that models both site-specific random intercepts and age-related random slopes (LME_INT+SLP_), (3) ComBat, and (4) ComBat with a generalized additive model (ComBat-GAM). Our test case for comparing harmonization methods was cortical thickness data aggregated from 29 sites, which included 1,343 cases with posttraumatic stress disorder (PTSD) (6.2-81.8 years old) and 2,067 trauma-exposed controls without PTSD (6.3-85.2 years old). We found that, compared to the other data harmonization methods, data processed with ComBat-GAM were more sensitive to the detection of significant case-control differences in regional cortical thickness (*X*^2^(3) = 34.339, *p* < 0.001), and case-control differences in age-related cortical thinning (*X*^2^(3) = 15.128, *p* = 0.002). Specifically, ComBat-GAM led to larger effect size estimates of cortical thickness reductions (corrected *p-values < 0.001*), smaller age-appropriate declines (corrected *p-values < 0.001*), and lower female to male contrast (corrected *p-values < 0.001*) in cases compared to controls relative to other harmonization methods. Harmonization with ComBat-GAM also led to greater estimates of age-related declines in cortical thickness (corrected *p-values < 0.001*) in both cases and controls compared to other harmonization methods. Our results support the use of ComBat-GAM for harmonizing cortical thickness data aggregated from multiple sites and scanners to minimize confounds and increase statistical power.

## Introduction

Large consortia, such as Enhancing Neuro Imaging Genetics through Meta-Analysis (ENIGMA) (Thompson et al., 2020), Cohorts for Heart and Aging Research in Genomic Epidemiology (CHARGE) (Hofer et al., 2020), and others have aggregated neuroimaging data acquired on many different scanners and recruited subjects at many different sites to conduct meta- and mega-analyses. By applying standardized analysis pipelines to extremely large datasets of thousands or tens of thousands of samples, consortia improve reliability, enhance reproducibility of results, amass sufficient statistical power to detect relatively small effect sizes, and support the ability to divide samples while retaining the power to delineate subsample (e.g., male vs female or young vs old) and interaction effects. The diverse ethnic, racial, geographic, and clinical demography of consortium data has provided results that are more representative of the wider population while also permitting exploration of clinical and neurobiological subtypes of neuropsychiatric disorders (Dennis et al., 2020; Thompson et al., 2020). Neuroimaging results generated by consortia are more robust and reproducible than studies that are generated by a single laboratory (Koshiyama et al., 2020), provided that consortia apply uniform methods to data originating from multiple sites and scanners.

However, several challenges are posed by the analysis of consortium data. A major concern of consortium-generated results is bias introduced by site-specific acquisition protocols and MRI scanners that may interact with site-specific demographic and clinical profiles (Radua et al., 2020).The challenge of *post hoc* combination of datasets stems partly from a lack of *a priori* harmonization of MRI acquisition sequences. Prospective data collection by consortia such as NCANDA (Brown et al., 2015), ABCD (Volkow et al., 2018), TRACK-TBI (Hicks et al., 2013), and others have prescribed harmonized acquisition parameters at study outset with the expectation of superior performance and obviating the need for post-acquisition harmonization. However, even prospective standardization and prescription of acquisition parameters results in significant variance attributed to sites for relatively short scan duration (e.g., 5 min) that can be reduced significantly by increasing scan duration (e.g., 25 min) (Noble et al., 2017). It remains unclear whether further post hoc harmonization of these datasets may improve sensitivity and power of analyses.

Various methods to harmonize neuroimaging data across sites are gaining acceptance and will become commonplace. However, there is little empirical evidence to support the use of a single method due to the lack of formal comparisons of available methods. In this study, we compared four harmonization methods. First, we tested linear mixed-effects modeling (LME), also known as the mixed-effects mega-analysis (ME-Mega) (Radua et al., 2020), with site as a random intercept (LME_INT_) to model the intercept location effects of site on brain measures. Second, we tested LME with both random intercept and age-related random slope for the site covariate (LME_INT+SLP_, see **Fig. 1**). Third, we used ComBat, a method originally developed to minimize batch effects present in data originating from multiple gene arrays (Johnson et al., 2007), and later adapted for neuroimaging data. ComBat is designed to remove site-associated differences while preserving variation due to biologically relevant variables such as age, sex, and diagnosis (Fortin et al., 2018). ComBat has been widely used to harmonize neuroimaging data including cortical thickness (Fortin et al., 2018), surface area, subcortical volumes (Radua et al., 2020), diffusion tensor imaging (Fortin et al., 2017; Hatton et al., 2020), and resting-state functional connectivity (Yu et al., 2018). Radua et al. (2020) reported that ComBat and LME_INT_ produced similar results when harmonizing cortical thickness, surface area, and subcortical volumes, while ComBat harmonization led to slightly higher statistical significance when performing between-group comparisons, in a multisite imaging study of schizophrenia. The fourth method, by Pomponio et al. (2020), improves on ComBat by modeling non-linear effects of age with a generalized additive model (GAM). ComBat-GAM allows for varied distributions of scale (multiplicative, or variance) and location (additive, or mean) effects, respectively.

**Figure 1.**
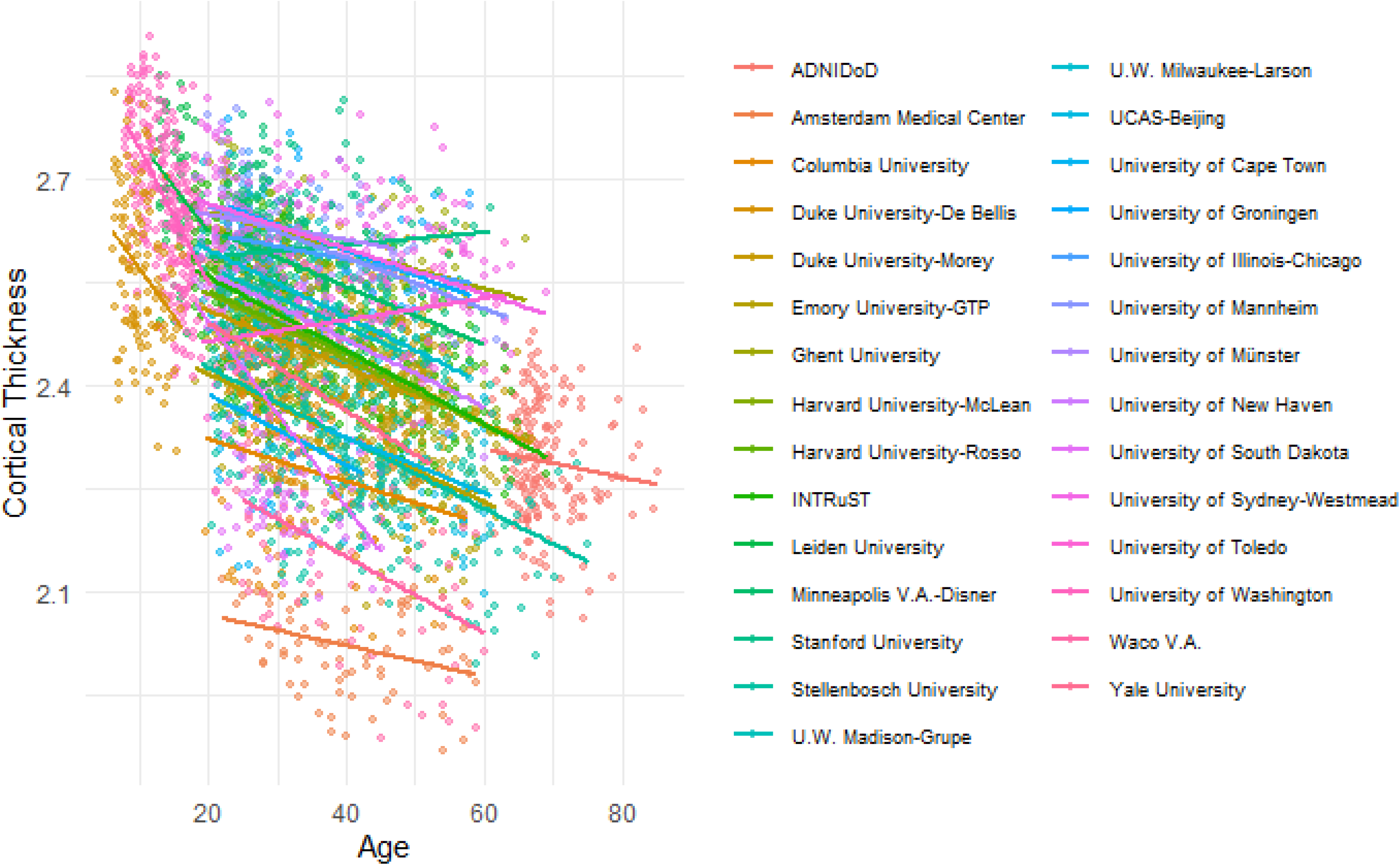
Scatter plots and age-related linear trends of mean cortical thickness averaged across regions for each study site. Participants are color-coded based on study site.

ComBat-GAM was designed to capture age-related non-linearities across the lifespan by fitting a GAM with a penalized nonlinear term. Pomponio et al. (2020) examined cortical and subcortical gray matter volumes without harmonization, harmonized by ComBat, and harmonized by ComBat-GAM in a large sample of 10,477 healthy subjects aggregated from 18 sites who ranged in age from 3-96 years. They reported that gray matter volumes harmonized by ComBat-GAM achieved the best performance in an age prediction task that minimized the difference between actual age and predicted age. They also found that ComBat-GAM, compared to other approaches, consistently led to improved prediction accuracy for each dataset in a leave-site-out validation experiment. However, Pomponio et al. (2020) only investigated data from healthy participants, which did not involve case-control comparisons, nor formal comparisons to LME methods.

Consequently, the goals of the present study were to investigate (1) the performance of ComBat-GAM for comparing clinical cases to controls, (2) how performance is influenced by age, and (3) how well performance characteristics compare to LME_INT_, LME_INT+SLP_, and ComBat. Although the random-effects meta-analysis (RE-Meta) has been widely used by ENIGMA projects (Zugman et al., 2020), we did not include RE-Meta in this study because several studies showed that LME and ComBat produce results with greater statistical power than RE- Meta (Boedhoe et al., 2017; Favre et al., 2019; Radua et al., 2020; van Rooij et al., 2018). The increase in power is based on the premise that the site effect being removed represents random noise, and its removal leads to larger effect sizes and greater efficiency requiring fewer subjects to reject the null hypothesis at a pre-specified power.

Data aggregated from 29 sites served as our test case for comparing harmonization methods. Subjects’ data was grouped into *cases* with PTSD (N=1,343) and trauma-exposed *controls* without PTSD (N=2,067). PTSD is associated with anatomical and functional alterations in widely distributed regions of the brain (Dennis et al., 2020; Logue et al., 2018; Wang et al., 2020). Military service members with PTSD and comorbid mild traumatic brain injury (mTBI) experience faster age-associated decline in cortical thickness than controls (Santhanam et al., 2019; Savjani et al., 2017). We hypothesized significant case-control differences in cortical thickness and age-related cortical thinning would be detectable in more brain regions by utilizing ComBat-GAM relative to LME_INT_, LME_INT+SLP_, and ComBat. We recognize that neither the method with the greatest number of regions reaching significance nor the method that maximizes the magnitude of regression coefficients reflects the true underlying cortical thickness - the so-called *ground truth* - which can only be measured definitively in post-mortem brains.

## Methods

### Participants

Data were obtained for secondary analysis from the ENIGMA-PGC PTSD Working Group. The dataset originated from 29 sites located on five continents (PTSD/Control N = 1,343/2,067). Demographic information is summarized in **Table 1**. Clinical measures and assessment of PTSD are explained in the Supplementary Materials. All study sites obtained approval from local institutional review boards or ethics committees. All participants provided written informed consent. Data is available upon request from the corresponding author.

**Table 1.**
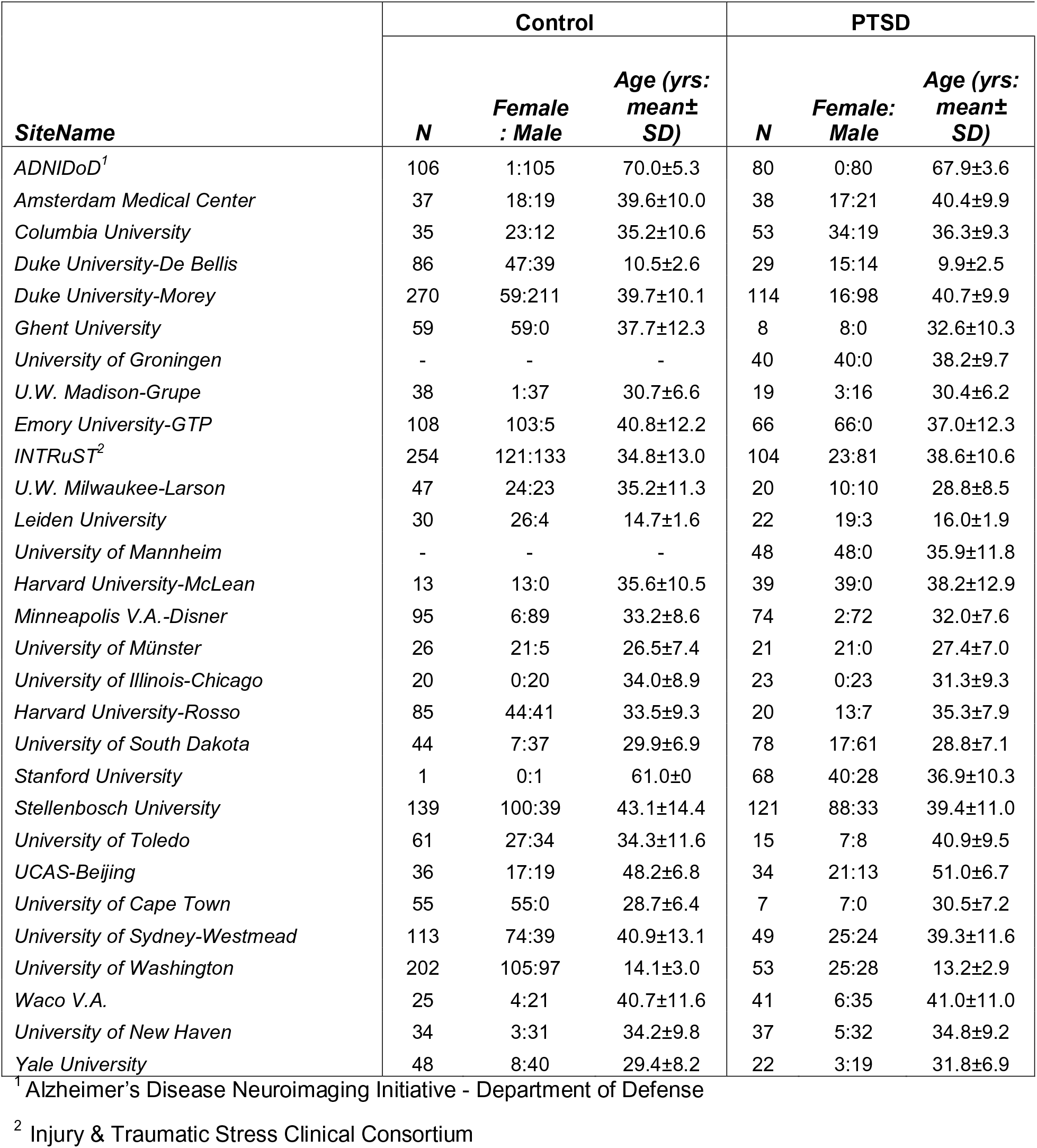
Demographic information per study site.

### Imaging Data Preprocessing

Anatomical brain images were preprocessed at Duke University through a standardized neuroimaging and QC pipeline developed by the ENIGMA Consortium (http://enigma.ini.usc.edu/protocols/imaging-protocols/) (Logue et al., 2018). Cortical thickness measurements were generated using the FreeSurfer software (https://surfer.nmr.mgh.harvard.edu) based on the Destrieux atlas (Destrieux et al., 2010) that contains 74 regions per hemisphere. Briefly, white matter surfaces were deformed toward the gray matter boundary at each surface vertex. Cortical thickness was calculated based on the average distance between the parcellated portions of white and pial surfaces within each region per participant. In each region, the cortical thickness of any participant more than 1.5 interquartile ranges (IQRs) below the first quartile or above the third quartile of the cortical thickness from all participants was defined as an outlier and was replaced by the mean cortical thickness averaged across participants of this region.

### ComBat Harmonization

ComBat removes the effects of site while preserving inherent biological variance in the data (Fortin et al., 2018). In the present study, PTSD diagnosis, age, and sex were designated as biological variables. The ComBat approach was implemented using R scripts (https://github.com/Jfortin1/ComBatHarmonization) running on RStudio (ver. 1.3.1073) and R (ver. 4.0.2). Unlike implementations of LME models that merge data harmonization and statistical analyses, ComBat and ComBat-GAM perform only harmonization and make harmonized data available to the user.

### ComBat-GAM Harmonization

PTSD diagnosis, age, and sex were designated as biological variables, and age was specified as a non-linear term in the model. The ComBat-GAM approach was implemented using Python (ver. 3.7.6) scripts (https://github.com/rpomponio/neuroHarmonize).

### Distribution of non-Harmonized, ComBat Harmonized, and ComBat-GAM Harmonized Data

The site-specific distribution of data harmonized by ComBat and ComBat-GAM was compared to non-harmonized data (data prior to harmonization). Using the R package *emmeans*, site-specific residuals (i.e., the absolute differences between the site-specific mean values and the mean value averaged across sites) and site-specific standard deviations for cortical thickness were compared across non-harmonized, ComBat harmonized, and ComBat-GAM harmonized data. The *p*-values were adjusted using the Tukey method for three pairwise comparisons (i.e., ComBat vs. non-harmonized, ComBat-GAM vs. non-harmonized, ComBat-GAM vs. ComBat).

### Statistical Models

In all models, we included sex, age, and PTSD diagnosis as fixed factors to estimate their effects on regional cortical thickness, and as covariates for testing interaction effects of interest. Either age by diagnosis interaction or sex by diagnosis interaction was included in the models as a fixed factor when the corresponding interaction was of interest. Linear modeling was used to analyze data harmonized by ComBat and data harmonized by ComBat-GAM. Cortical thickness data without harmonization was entered into the LME models. The LME_INT_ models employed study site as a random factor to model random intercepts. The LME_INT+SLP_ modeled both the site-specific random intercepts and age-related random slopes to reflect different age-related slopes in cortical thickness across sites (**Fig. 1**). The Bonferroni method was employed to correct for multiple testing of 148 cortical regions with a corrected α = 0.0003 (0.05/148). The R packages *lme4,* and *lmerTest were* used to calculate regression coefficients and statistical significance for the random effects models.

The number of regions with significant findings and the magnitude of effect size was compared separately between the 4 harmonization methods. A chi-squared test was used to compare the number of cortical regions showing significant effects. The region-specific regression coefficients were compared using repeated-measures ANOVA from the R package *afex*. If the omnibus ANOVA results were statistically significant, then post-hoc pairwise comparisons of the 4 harmonization methods were conducted using the R package *emmeans*. The *p*-values were adjusted using the Tukey method for the 6 pairwise comparisons made with the outputs of the 4 harmonization methods.

## Results

### Distribution of non-Harmonized, ComBat Harmonized, and ComBat-GAM Harmonized Data

Residuals represent the absolute value of the difference between the mean cortical thickness value of subjects at a given site and the mean cortical thickness value of all sites. Relative to non-harmonized data (**Fig. 2)**, the cortical thickness data harmonized by ComBat (controls: *t-values* = -10.162∼-2.908, *p-values* = <0.001∼0.014 corrected; PTSD: *t-values* = -10.028∼-2.865, *p-values* = <0.001∼0.016 corrected; across regions) and ComBat-GAM (controls: *t-values* = -10.150∼-1.601, *p-values* = <0.001∼0.254 corrected; PTSD: *t-values* = -9.978∼-1.907, *p-values* = <0.001∼0.146 corrected; across regions) resulted in smaller residuals. There was no significant difference in residuals between ComBat and ComBat-GAM harmonized data (controls: *t-values* = -1.645∼0.197, *p-values* = 0.999 corrected; PTSD: *t-values* = -1.662∼0.173, *p-values* = 0.999 corrected; across regions).

**Figure 2.**
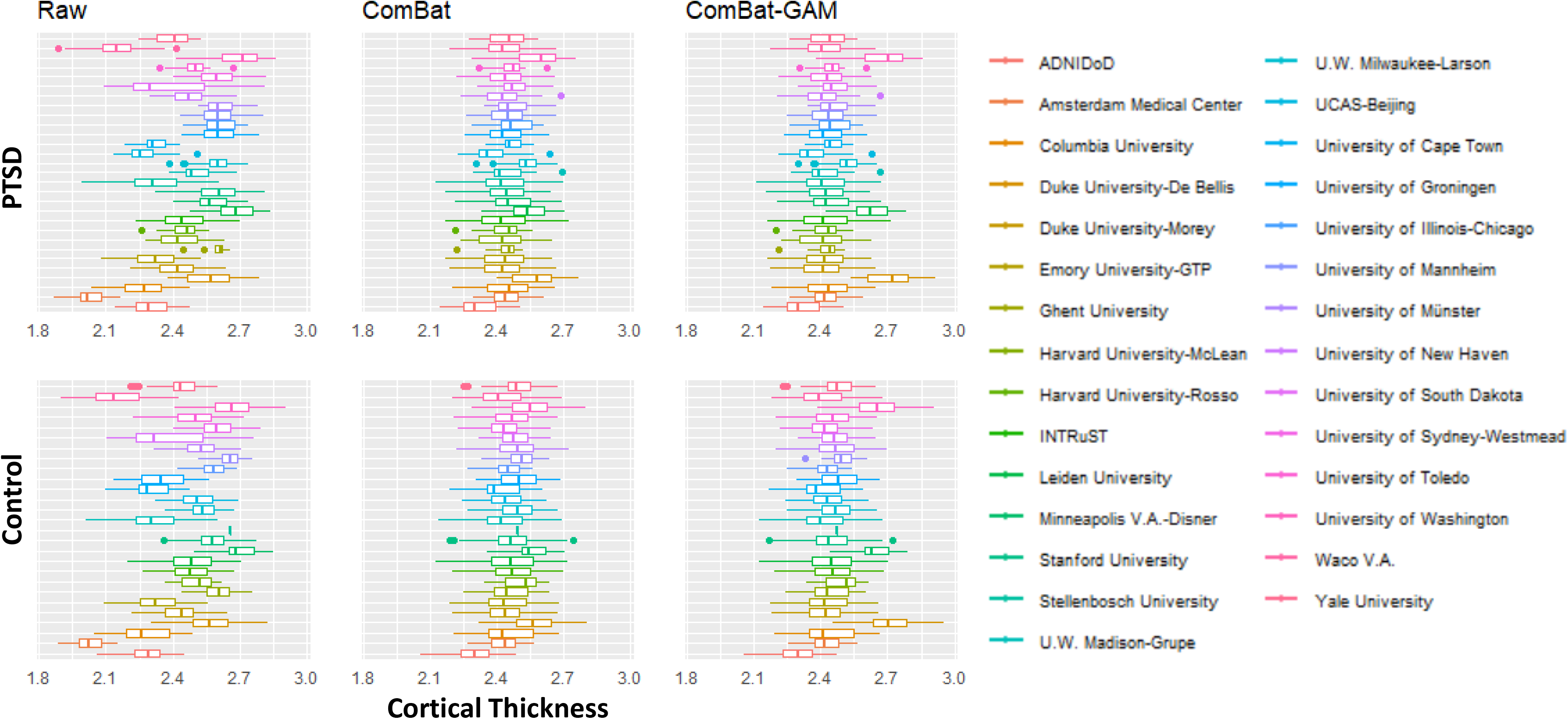
Site-specific cortical thickness averaged across regions for non-harmonized, ComBat harmonized, and ComBat-GAM harmonized data in trauma-exposed participants with and without PTSD.

There was no significant difference in the standard deviations between all data-pairings across all regions (controls: *t-values* = -0.576∼2.584, *p-values* = 0.788∼2.584 corrected; PTSD: *t-values* = -0.545∼3.070, *p-values* = 0.654∼1.000 corrected). These results suggest that both ComBat and ComBat-GAM efficiently reduce differences in site-specific intercepts, but do not change differences in site-specific variances. The age-related distribution of non-harmonized, ComBat harmonized, and ComBat-GAM harmonized data are shown in **Fig. 3**.

**Figure 3.**
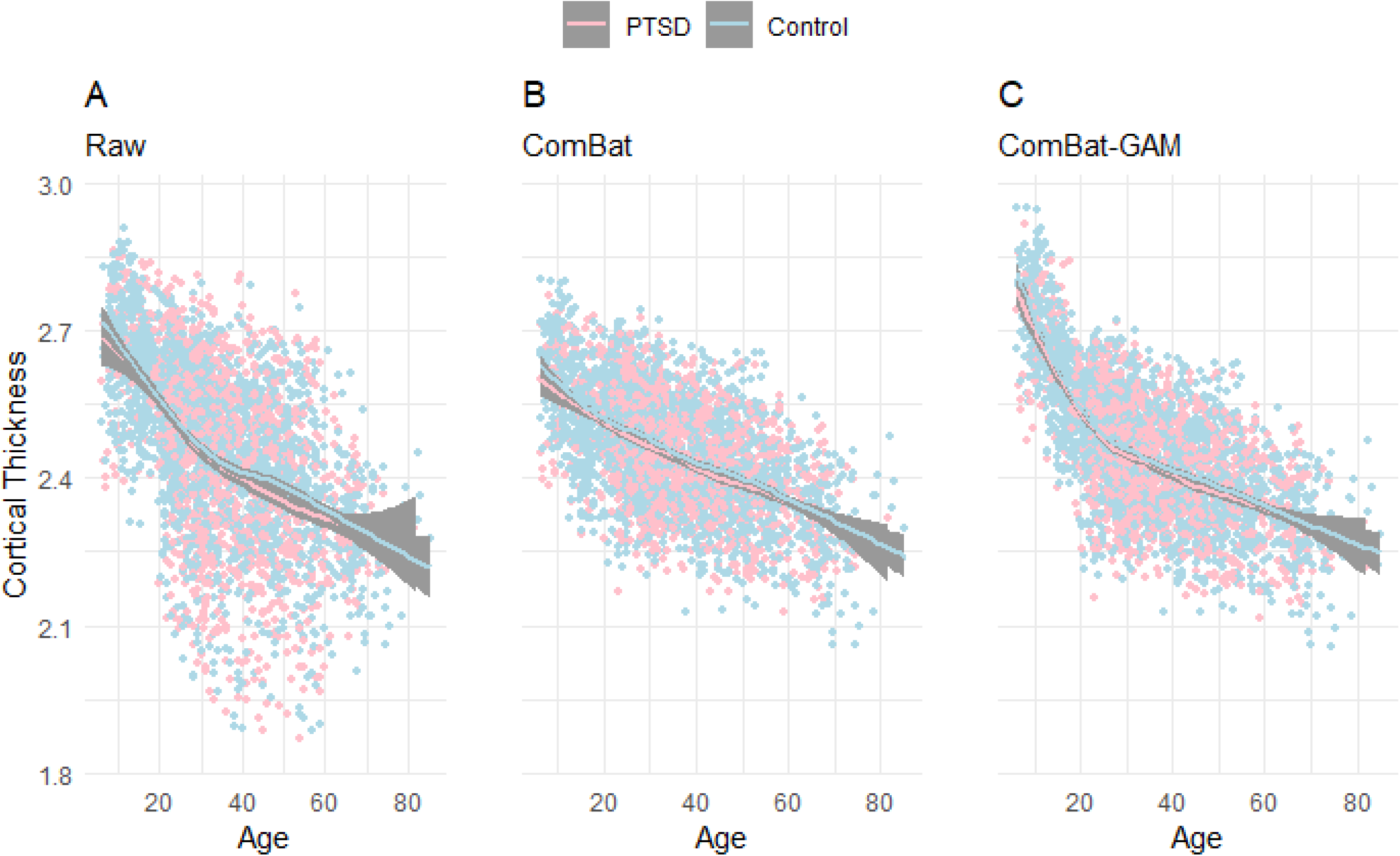
Scatter plots and non-linear trends of mean cortical thickness averaged across regions for (A) non-harmonized data, (B) ComBat harmonized data, and (C) ComBat-GAM harmonized data, in trauma-exposed participants with and without PTSD.

### Main effects of Age

As shown in **Fig. 4A&B**, the number of regions showing a significant main effect of age was significantly different across harmonization methods (*X*^2^(3) = 223.550, *p* < 0.001). The age-related declines in cortical thickness were detected by ComBat-GAM and ComBat in 148 (100%) regions, by LME_INT_ in 145 (98.0%) regions, and by LME_INT+SLP_ in 78 (52.7%) regions. It seems that LME_INT+SLP_ harmonization was less efficient in detecting age-related differences in cortical thickness.

**Figure 4.**
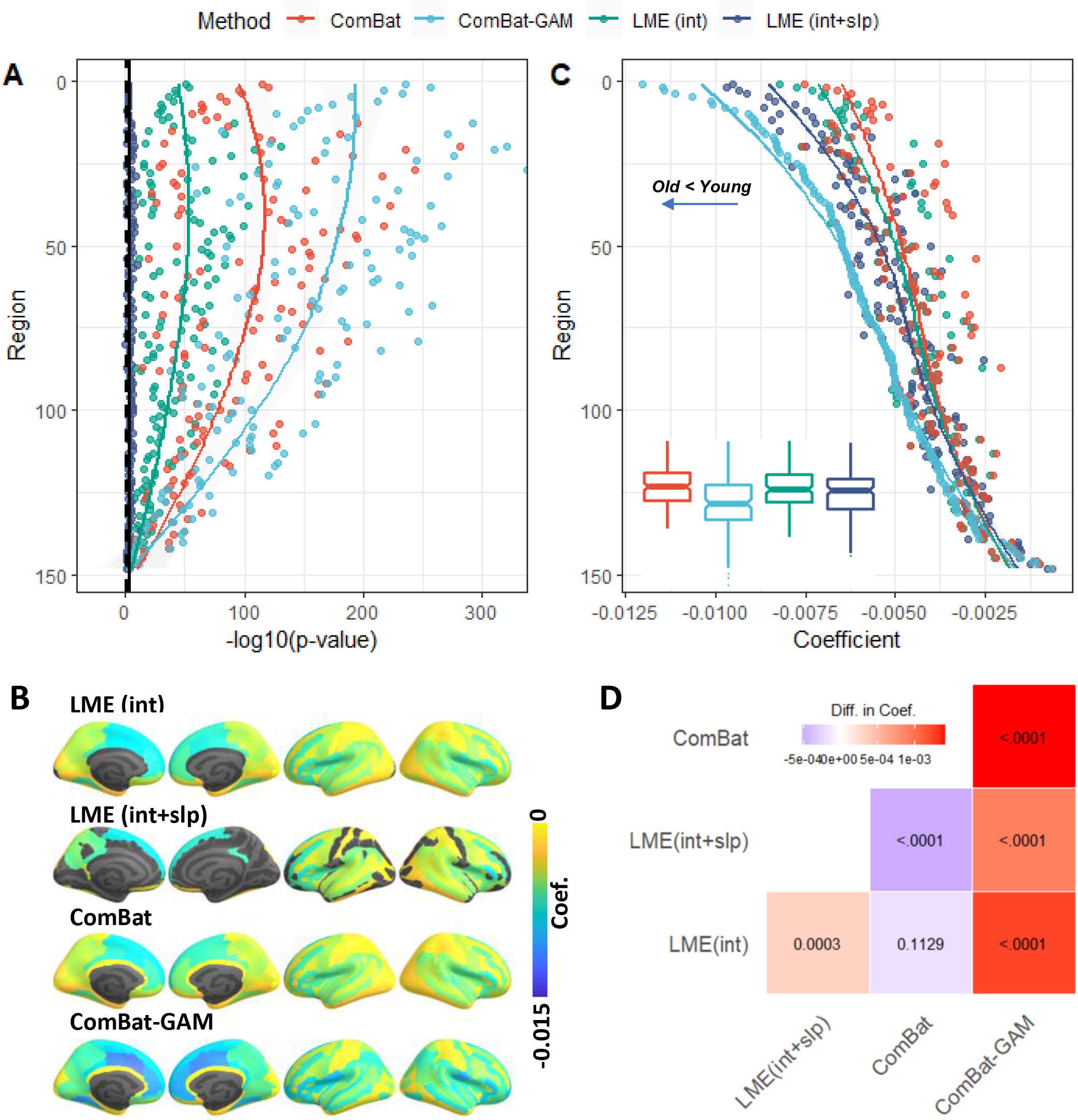
Main effect of age. (A) negative log-transformed statistical significance, i.e. –log_10_(p). The dashed and solid vertical lines represent thresholds *p* = .05 (uncorrected) and *p* = .05 (Bonferroni corrected), respectively. (B) Regions show a significant main effect of age. The color bar represents the magnitude of the regression coefficient. Cooler colors represent age-related declines in cortical thickness. (C) Magnitude of regression coefficients. The ordering of regions from top to bottom in both (A) and (C) is by ascending order of regression coefficients from cortical thickness data harmonized by ComBat-GAM. Boxplots of the regression coefficients per harmonization methods were also displayed. (D) Corrected p-values associated with pairwise comparisons of regression coefficients. The color bar represents differences in regression coefficients between row-labeled and column-labeled harmonization methods. LME_INT_, LME models site-specific random intercept. LME_INT+SLP_, LME models both site-specific random intercepts and age-related random slopes.

We found that 53.8%, 100%, and 100% of regions detected by LME_INT_ were identified by LME_INT+SLP_, ComBat, and ComBat-GAM, respectively. All regions detected by LME_INT+SLP_ were identified by the other three methods. 98.0%, 52.7%, and 100% of regions detected by ComBat were also identified by LME_INT_, LME_INT+SLP_, and ComBat-GAM, respectively. 98.0%, 52.7%, and 100% of regions detected by ComBat-GAM were also identified by LME_INT_, LME_INT+SLP_, and ComBat, respectively.

As shown in **Fig. 4C**, the regression coefficients were significantly different across harmonization methods (*F*(1.4, 212.3) = 123.25, *p* < 0.001). Post-hoc analyses (**Fig. 4D**) showed that all the other three methods produced higher age-related regression coefficients (i.e., lower estimates of age-related declines) than ComBat-GAM, while LME_INT+SLP_ produced lower age-related regression coefficients than LME_INT_ and ComBat.

### Main effects of diagnosis

As shown in **Fig. 5A&B**, the number of regions showing a significant main effect of diagnosis was significantly different across harmonization approaches (*X*^2^(3) = 34.339, *p* < 0.001). Case-related reductions in cortical thickness were found by ComBat-GAM in 25 (16.7%) regions, by ComBat in 6 (4.0%) regions, by LME_INT_ in 4 (2.7%) regions, and by LME_INT+SLP_ in 4 (2.7%) regions. The regions discovered by ComBat-GAM include those within the salience network (SN; right anterior cingulate cortex and bilateral insula regions), executive control network (ECN; left intraparietal sulcus and bilateral supramarginal gyri), default mode network (DMN; bilateral ventromedial prefrontal cortex, and bilateral precuneus), and bilateral superior and inferior temporal gyri and sulci, which are consistent with previous reports (Shalev et al., 2017).

**Figure 5.**
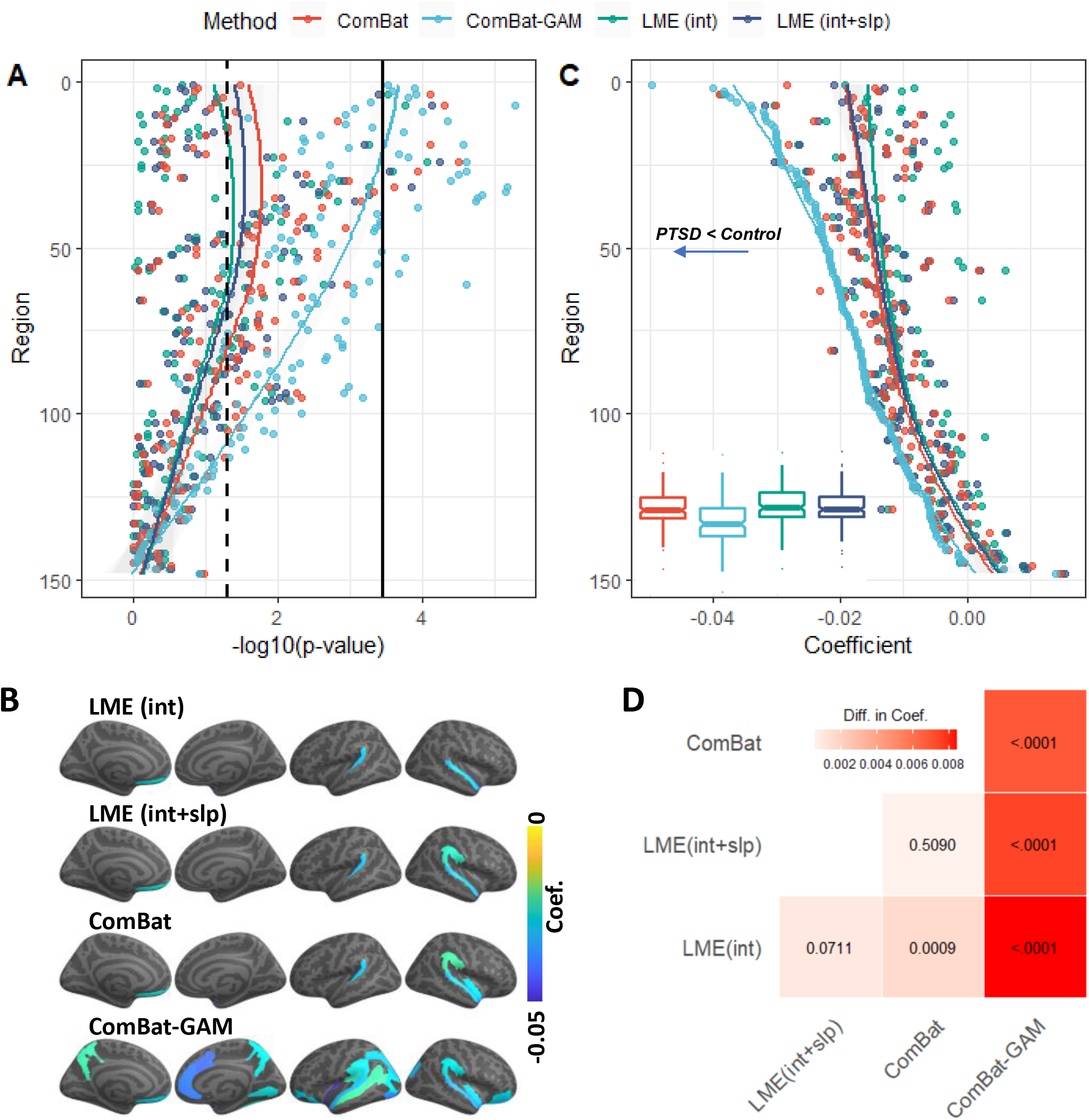
Case-control main effects. (A) negative log-transformed statistical significance, i.e. – log_10_(p). The dashed and solid vertical lines represent thresholds *p* = .05 (uncorrected) and *p* = .05 (Bonferroni corrected), respectively. (B) Regions that show a significant case-control main effect. The color bar represents the magnitude of the regression coefficient. Cooler colors mean lower cortical thickness in PTSD than controls. (C) Magnitude of regression coefficients. The ordering of regions from top to bottom in both (A) and (C) is by ascending order of regression coefficients from cortical thickness data harmonized by ComBat-GAM. Boxplots of the regression coefficients per harmonization methods were also displayed. (D) Corrected p-values associated with pairwise comparisons of regression coefficients. The color bar represents the differences of regression coefficients between row-labeled and column-labeled harmonization approaches. LME_INT_, LME models site-specific random intercept. LME_INT+SLP_, LME models both site-specific random intercepts and age-related random slopes.

We found that 75.0%, 75.0%, and 100% of regions detected by LME_INT_ were identified by LME_INT+SLP_, ComBat, and ComBat-GAM, respectively. 75.0%, 75.0%, and 100% of regions detected by LME_INT+SLP_ were identified by LME_INT_, ComBat, and ComBat-GAM, respectively 66.7%, 66.7%, and 100% of regions detected by ComBat were identified by LME_INT_, LME_INT+SLP_, and ComBat-GAM, respectively. 16.0%, 16.0%, and 24.0% of regions detected by ComBat-GAM were identified by LME_INT_, LME_INT+SLP_, and ComBat, respectively. Importantly, data harmonized with ComBat-GAM led to the detection of all regions identified by the other three methods.

As shown in **Fig. 5C**, regression coefficients were different across harmonization methods (*F*(1.6, 163.0) = 169.55, *p* < 0.001). Post-hoc analyses (**Fig. 5D**) showed that the three other methods produced higher case-related regression coefficients (i.e., lower estimates of case-related cortical thickness reduction) than ComBat-GAM, while LME_INT_ produced higher regression coefficients than ComBat.

### Age by Diagnosis Interaction

As shown in **Fig. 6A, B**, significant age by diagnosis interactions were detected by ComBat-GAM in 5 regions, while no significant interactions were detected by ComBat, LME_INT_, and LME_INT+SLP_. ComBat-GAM outperformed the other methods in detecting this interaction effect (*X*^2^(3) = 15.128, *p* = 0.002). Age-related declines in cortical thickness were slower in cases than controls for 5 regions within the DMN, which include the left middle-posterior part of the cingulate gyrus and sulcus, the right marginal branch of the cingulate sulcus, the right superior frontal sulcus in ECN, right inferior temporal areas that include the right medial occipital-temporal sulcus and lingual sulcus, and the right fusiform gyrus. The linear (**Fig. S1**) and non-linear (**Fig. S2**) fits of the age-related distributions of cortical thickness in the 5 regions were shown in supplementary.

**Figure 6.**
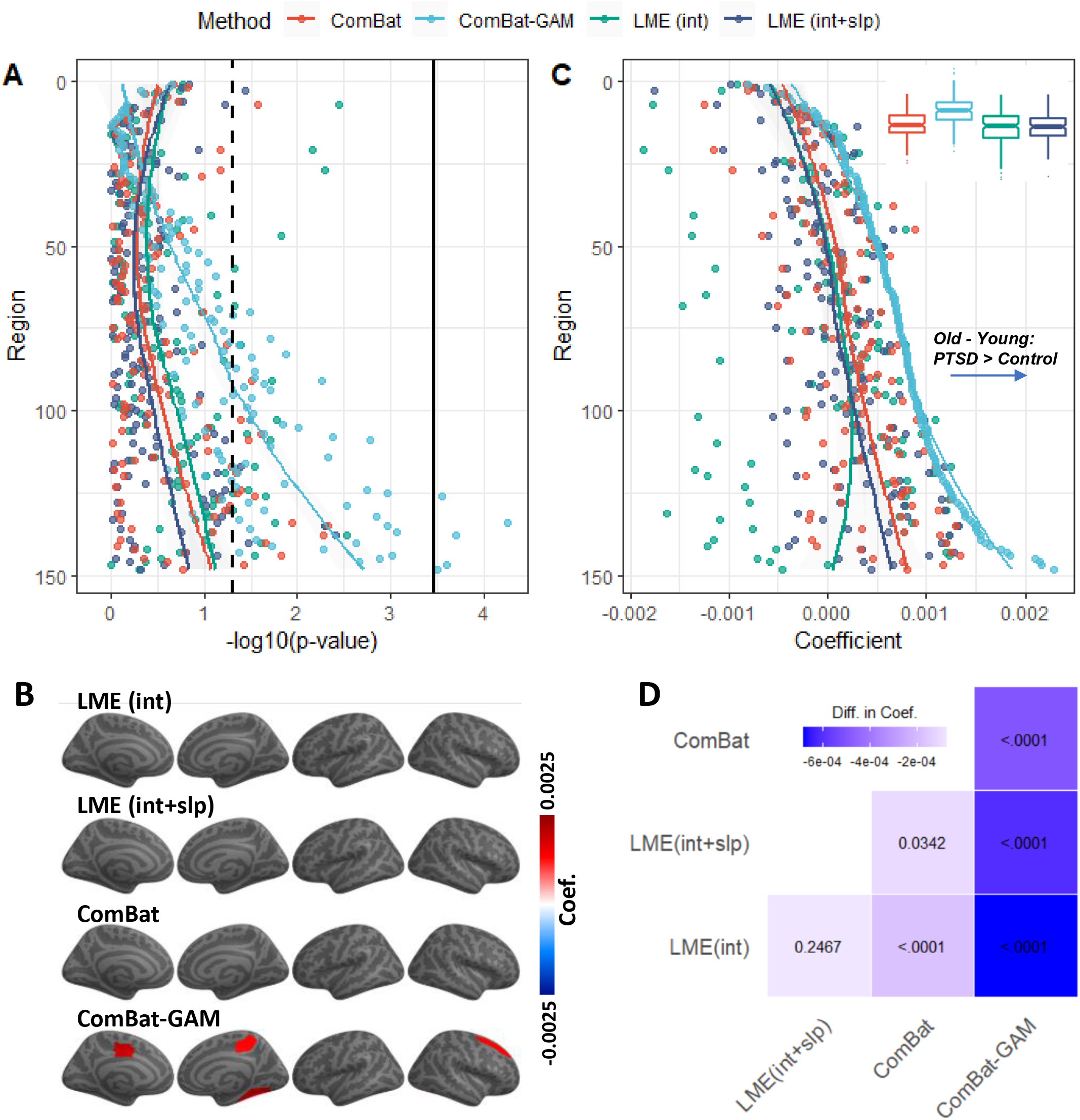
Interaction of age and diagnosis. (A) negative log-transformed statistical significance, i.e. –log_10_(p). The dashed and solid vertical lines represent thresholds *p* = .05 (uncorrected) and *p* = .05 (Bonferroni corrected), respectively. (B) Regions show significant age by diagnosis interaction. The color bar represents the magnitude of the regression coefficient. Warmer colors mean that age-related declines in cortical thickness are slower in PTSD than controls. (C) Magnitude of regression coefficients. The ordering of regions from top to bottom in both (A) and (C) is by ascending order of regression coefficients from cortical thickness data harmonized by ComBat-GAM. Boxplots of the regression coefficients per harmonization methods were also displayed. (D) Corrected p-values associated with pairwise comparisons of regression coefficients. The color bar represents differences in regression coefficients between row-labeled and column-labeled harmonization methods. LME_INT_, LME models site-specific random intercept. LME_INT+SLP_, LME models both site-specific random intercepts and age-related random slopes.

As shown in **Fig. 6C**, regression coefficients differed across harmonization methods (*F*(1.2, 184.0) = 121.99, *p* < 0.001). Post-hoc analyses (**Fig. 6D**) showed that the three other methods compared to ComBat-GAM produced lower age-related regression coefficients in cases than controls (i.e., higher estimates of age-related declines in cortical thickness in cases than controls), while both LME_INT_ and LME_INT+SLP_ produced lower regression coefficients than ComBat.

### Main effects of Sex

As shown in **Fig. 7A, B**, the number of regions showing a significant main effect of sex was not significantly different across harmonization methods (*X*^2^(3) = 2.749, *p* = 0.432).

**Figure 7.**
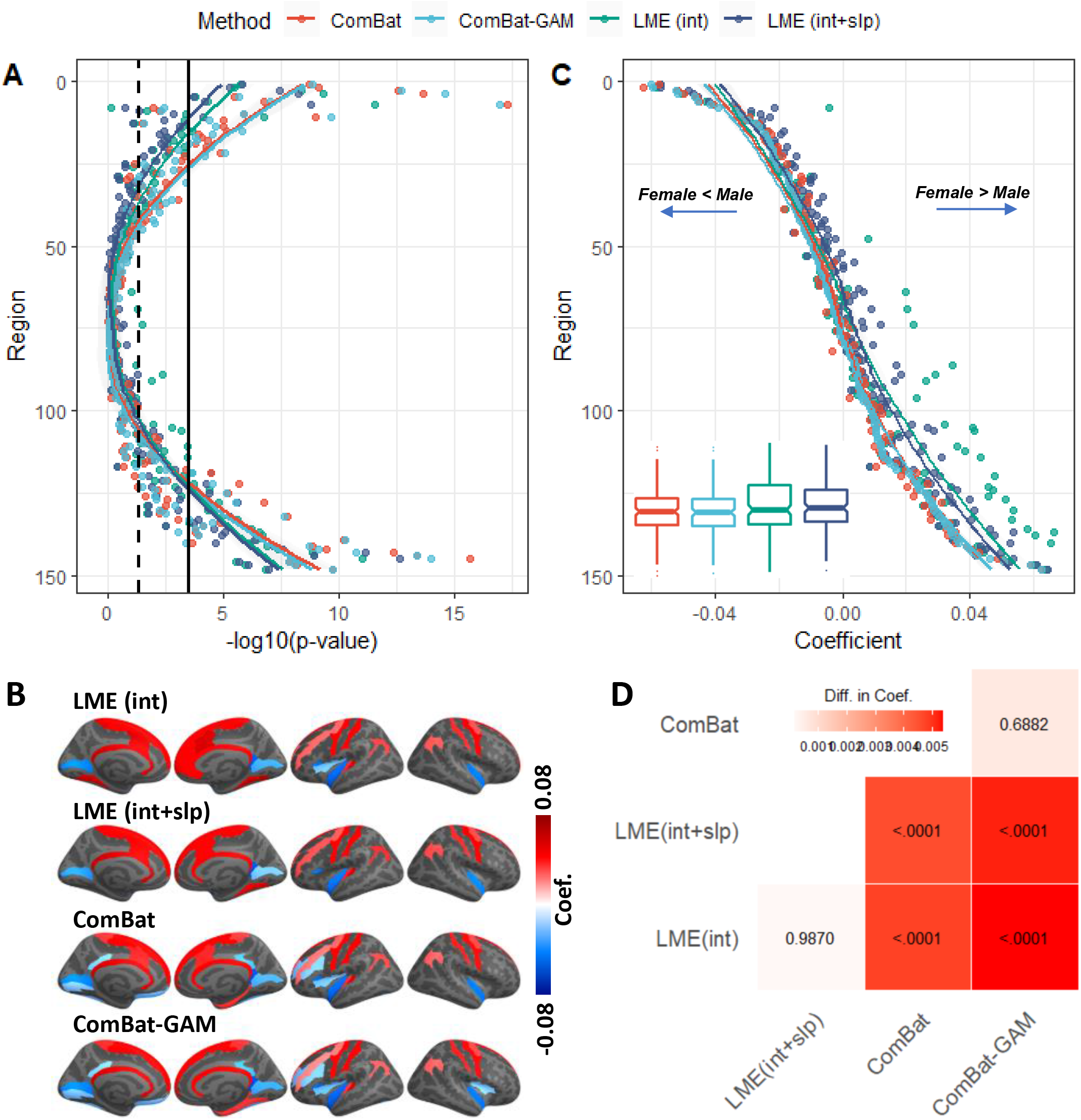
Main effects of sex. (A) negative log-transformed statistical significance, i.e. –log_10_(p). The dashed and solid vertical lines represent thresholds *p* = .05 (uncorrected) and *p* = .05 (Bonferroni corrected), respectively. (B) Regions show a significant main effect of sex. The color bar represents the magnitude of the regression coefficient. Cooler (warmer) colors indicate lower (higher) cortical thickness in females compared to males. (C) Magnitude of regression coefficients. The ordering of regions from top to bottom in both (A) and (C) is by ascending order of regression coefficients from cortical thickness data harmonized by ComBat-GAM. Boxplots of the regression coefficients per harmonization methods were also displayed. (D) Corrected p-values associated with pairwise comparisons of regression coefficients. The color bar represents the difference in regression coefficients between row-labeled and column- labeled harmonization approaches. LME_INT_, LME models site-specific random intercept. LME_INT+SLP_, LME models both site-specific random intercepts and age-related random slopes.

Differences between males and females in cortical thickness were detected by ComBat-GAM and ComBat in 38 regions, by LME_INT_ in 32 regions, and by LME_INT+SLP_ in 28 regions. The analyses based on ComBat-GAM harmonization showed that females had greater cortical thickness than males in bilateral precentral and postcentral regions, bilateral anterior cingulate cortex, bilateral superior frontal gyri, bilateral angular gyri, bilateral medial occipito-temporal sulci and lingual sulci, left frontal pole, left superior temporal sulci, and right parahippocampal gyrus. By contrast, males had greater cortical thickness than females in bilateral inferior temporal regions, left rectus, left planum polare of the superior temporal gyrus, left vertical ramus of the anterior segment of the lateral sulcus, bilateral calcarine sulci, left insula, left inferior and middle frontal sulci, left orbital sulci, right ventral posterior cingulate cortex, right temporal pole.

We found that 87.5%, 87.5%, and 84.4% of regions detected by LME_INT_ were also identified by LME_INT+SLP_, ComBat, and ComBat-GAM, respectively. We found that 100%, 96.4%, and 92.9% of regions detected by LME_INT+SLP_ were also identified by LME_INT_, ComBat, and ComBat-GAM, respectively. We found that 73.7%, 71.1%, and 94.7% of regions detected by ComBat were also identified by LME_INT_, LME_INT+SLP_, and ComBat-GAM, respectively. We found that 71.1%, 68.4%, and 94.7% of regions detected by ComBat-GAM were also identified by LME_INT_, LME_INT+SLP_, and ComBat, respectively.

As shown in **Fig. 7C**, the regression coefficients were different across harmonization methods (*F*(1.2, 173.6) = 54.06, *p* < 0.001). Post-hoc analyses (**Fig. 7D**) showed that both LME_INT_ and LME_INT+SLP_ produced higher regression coefficients (i.e., higher estimates of cortical thickness in females than males) than ComBat and ComBat-GAM.

### Sex by PTSD diagnosis interaction

As shown in **Fig. 8A, B**, no significant sex by diagnosis interactions were found using data from the 4 the harmonization methods. As shown in **Fig. 8C**, the regression coefficients were significantly different across harmonization approaches (*F*(1.2, 184.0) = 121.99, *p* < 0.001). Post-hoc analyses (**Fig. 8D**) showed the three other methods produced higher regression coefficients (i.e., higher estimates of the female-male contrast of cortical thickness in cases than controls) than ComBat-GAM, while LME_INT_ produced higher regression coefficients than the three other methods.

**Figure 8.**
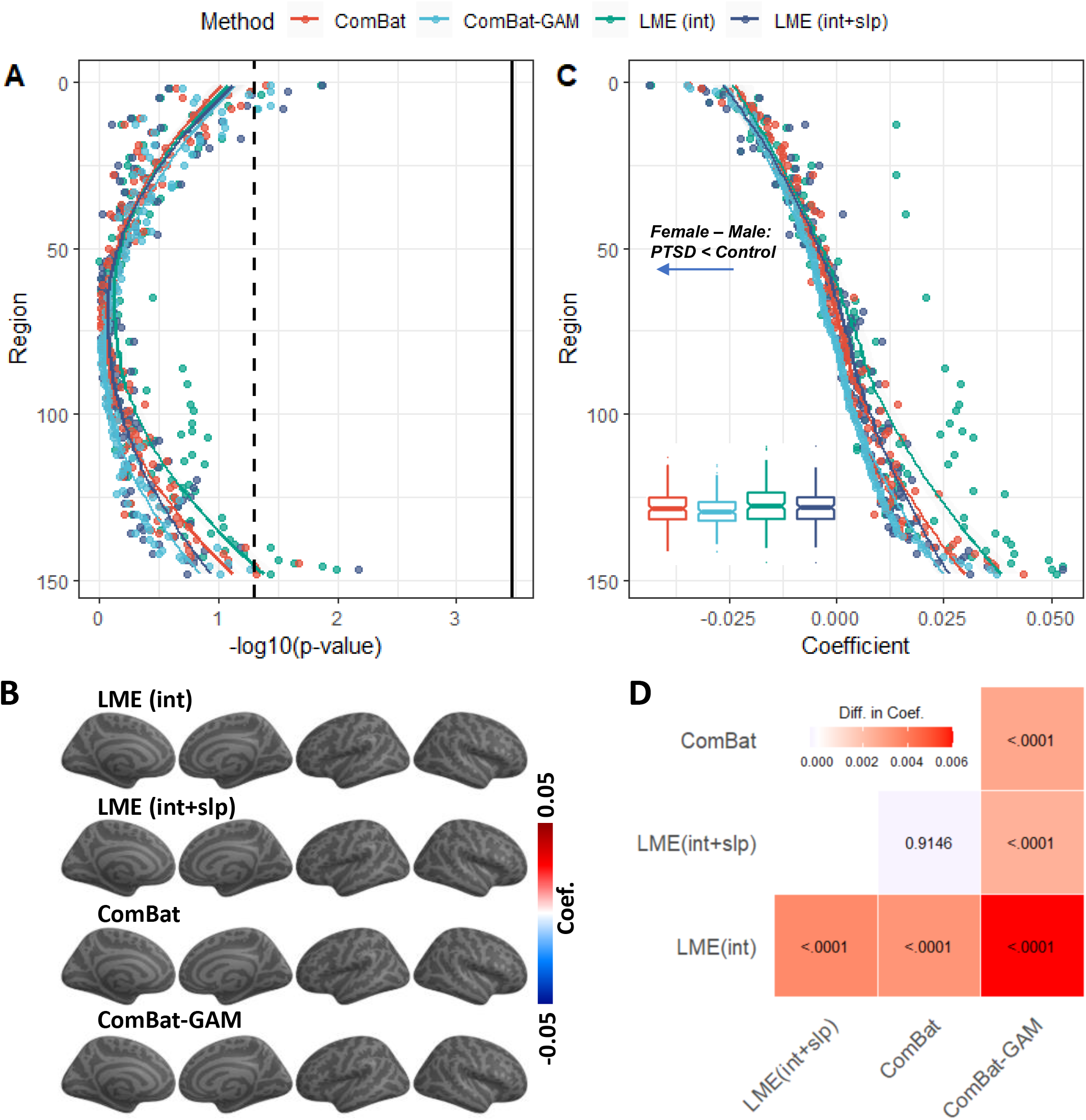
Sex by diagnosis interaction. (A) negative log-transformed statistical significance, i.e. –log_10_(p). The dashed and solid vertical lines represent thresholds *p* = .05 (uncorrected) and *p* = .05 (Bonferroni corrected), respectively. (B) No region shows significant sex by diagnosis interaction. The color bar represents the magnitude of the regression coefficient. (C) The magnitude of regression coefficients. The ordering of regions from top to bottom in both (A) and (C) is by ascending order of regression coefficients from cortical thickness data harmonized ComBat-GAM. Boxplots of the regression coefficients per harmonization methods were also displayed. (D) Corrected p-values associated with pairwise comparisons of regression coefficients. The color bar represents the difference in regression coefficients between row- labeled and column-labeled harmonization methods. LME_INT_, LME models site-specific random intercept. LME_INT+SLP_, LME models both site-specific random intercepts and age-related random slopes.

## Discussion

We compared the performance of four harmonization methods by applying them to cortical thickness data in participants grouped into clinical cases and controls from 29 different sites. The four harmonization methods included LME_INT_, LME_INT+SLP_, ComBat, and ComBat-GAM. We acknowledge at the outset the number of regions reaching significance by any given method does not necessarily reflect the *ground truth*. ComBat and ComBat-GAM harmonization detected the greatest number of regions with significant age-related declines in cortical thickness, LME_INT+SLP_ detected the fewest number of significant regions, while LME_INT_ detected an intermediate number of regions. Consistent with our *a priori* hypothesis, data harmonized with ComBat-GAM, relative to the other harmonization methods, led to the detection of more regions with significant case-related reductions in cortical thickness, and more regions displaying slower rates of age-related cortical thinning in cases than controls. There were no significant differences between the 4 harmonization methods in detecting sex-related differences in cortical thickness. A comparison of the regression coefficients showed that ComBat-GAM, relative to the other methods, produced higher estimates of cortical thickness reduction, lower estimates of age-related cortical thickness decline, and lower female-male contrast estimates in cases compared to controls. ComBat-GAM showed higher estimates for age-related declines in cortical thickness in cases and controls as compared to other harmonization methods. Both ComBat and ComBat-GAM methods produced lower cortical thickness estimates in females than males when compared to LME_INT_ and LME_INT+SLP_.

ComBat models the expected values of the imaging features as a linear combination of the biological variables and the site effects whose error term is modulated by additional site-specific scaling factors (Fortin et al., 2018). It also uses empirical Bayes to improve the estimation of the model parameters in studies with small sample size. (Radua et al., 2020) used cortical thickness, surface area, and subcortical volume data in cases and controls from ENIGMA-Schizophrenia to compare ComBat to random-effects meta-analysis and random-effects mega-analysis, which we term LME_INT_ in the present study. They reported that ComBat delivered more results that were statistically significant than random-effects meta-analyses, and slightly more than LME_INT_. However, they did not report results of non-linear age effects on cortical thickness, which are well documented (Frangou et al., 2021; Pomponio et al., 2020; Walhovd et al., 2017), nor did they report on effects of group membership on age-related changes in cortical thickness. By contrast, Pomponio et al. (2020) developed ComBat-GAM to support harmonization of neuroimaging data with age-related non-linearities or other variables by investigating cortical and subcortical gray matter volumes in 10,477 healthy subjects from 18 sites ranging in age from 3-96 years. They concluded that ComBat-GAM is superior to ComBat at predicting age based on regional volume data. However, Pomponio et al. (2020) only investigated healthy participants, which lacked guidance on harmonization of data used to make case-control comparisons. Moreover, prior studies did not report the magnitude of regression coefficients obtained from various harmonization methods, in spite of an urgent plea by researchers to understand how harmonization methodology influences the output of statistical models run on harmonized data.

Our study sought to fill these gaps by formally comparing regression coefficients and the number of regions showing statistically significant results, including the case-control differences in cortical thickness across the lifespan. Harmonization with ComBat-GAM was the most efficient at detecting case-control differences as evidenced by significantly more regional findings as compared to other harmonization methods. ComBat-GAM was also one of the most efficient methods at detecting age-effects in cortical thickness, and the only method to uncover regions with different rates of age-related cortical thinning in cases compared to controls.

Whereas we have no collateral information to corroborate the findings from ComBat-GAM harmonization pertaining to case-control differences or age-dependent case-control differences, we have reliable evidence of age-related patterns of cortical thickness across the lifespan (Frangou et al., 2021; Mutlu et al., 2013). One caveat is that motion related artifacts, which are associated with lower cortical thickness measurements, increase with age (Savalia et al., 2017). Consequently, reduced cortical thickness with aging may be partially explained by motion-induced reduction in apparent cortical thickness. Nonetheless, **Fig. 3B** shows concrete evidence of erroneous harmonization by ComBat that is handled correctly in **Fig. 3C** by ComBat-GAM as corroborated by independent studies, which demonstrate that the highest cortical thickness occurs in childhood and that age is negatively correlated to cortical thickness with a steeper slope up to the third decade of life then more gradual thereafter (Frangou et al 2021; Mutlu et al 2013). By contrast, ComBat harmonized data along a linear pattern with age throughout the lifespan. Thus, ComBat-GAM harmonization may be advantageous, particularly for consortia studies of participants of all ages.

The performance of ComBat-GAM is attributable to its algorithm. LME models assume that the error terms follow the same normal distribution at all sites, which is rarely the case (Radua et al., 2020). ComBat overcomes this shortcoming by assuming different normal distributions at different sites for the error terms (Radua et al., 2020). ComBat-GAM further improves on ComBat by assuming a normal distribution as the prior for the intercept location effect and an inverse-gamma distribution as the prior for the scale effect of the sites. It also uses the generalized additive models (GAMs) to capture the non-linear variations in age-related changes in cortical thickness while avoiding overfitting (Pomponio et al., 2020). In this study, ComBat-GAM appears to be the best method to capture the cortical variations associated with age, and therefore enhance its ability to accurately capture variations associated with diagnosis and sex. By contrast, harmonization with LME_INT+SLP_ produced the smallest number of regions with age-related changes. LME_INT+SLP_ allows the relationship between age and cortical thickness to be different across sites, which may reduce the variations of cortical thickness that could be explained by the fixed effect of age.

We found that the age-related decline in cortical thickness was slower in cases than controls for 5 regions, but only for data harmonized with ComBat-GAM. As shown in supplementary **Fig. S1** and **S2**, cases compared to controls exhibited lower cortical thickness in youth and higher cortical thickness in elderly in the 5 regions. It is possible that PTSD induces more powerful cortical thinning in youth and delayed age-appropriate declines in cortical thickness in elderly. This explanation is partly consistent with previous findings that maltreated youth with versus without chronic PTSD have smaller volumes in the right ventromedial prefrontal cortex (Morey et al., 2016) and posterior brain structures (De Bellis et al., 2015). More studies are warranted to test whether case-control differences in age-related cortical thinning is overfit by ComBat-GAM.

A study by Ritchie et al. (2018) that examined sex-differences in adults from UK Biobank (2,750 females; males 2,466; 45-80 years old) found greater cortical thickness in the postcentral, superior, and inferior parietal, and supramarginal gyri in females, whereas males had greater cortical thickness in the ventromedial prefrontal cortex and rostral anterior cingulate cortex. Similarly, in our study, data harmonization with ComBat-GAM showed that females have greater cortical thickness in prefrontal cortex, inferior parietal regions, cingulate cortex, and left temporal pole, whereas males had greater cortical thickness in ventromedial prefrontal cortex, bilateral insula, posterior cingulate areas, and occipital lobe. We found no statistically significant difference between harmonization methods in the number of regions showing sex-effects.

These results suggest that all four harmonization methods effectively detect regions with sex-related changes in cortical thickness. While we did not formally test harmonization methods to detect age-related sex differences in cortical thickness, Frangou et al. (2021) report that sex-differences in cortical thickness are age dependent.

The comparison of the regression coefficients showed that the selection of harmonization methods may overestimate or underestimate effects of interest, even though the corresponding comparisons of the number of regions exhibiting significant effects were identical between methods. ComBat-GAM, relative to the other methods, generated higher estimates of reductions in cortical thickness, and lower estimates of age-appropriate declines as well as female to male contrast in cases compared to controls. ComBat-GAM also generated higher estimates of age-related declines in cortical thickness in both cases and controls. Both ComBat and ComBat-GAM estimated lower cortical thickness in females than males when compared to LME_INT_ and LME_INT+SLP_. This knowledge is critical to interpreting statistical outputs. For instance, the magnitude of reductions in cortical thickness per year are biased by the harmonization method being used.

In reporting that ComBat-GAM is more sensitive than other methods, we must be clear to specify our narrow definition of “sensitive”, as the harmonization method that leads to the maximum number of brain regions with statistically significant effects. In fact, this metric does not necessarily determine better performance if we adopt a preferred definition, namely the method that produces results that are most consistent with the *ground truth*. Unfortunately, identifying ground truth is a challenging proposition, but we consider two options that may be informative and feasible. The first option is to acquire MRI scans and calculate cortical thickness on a variety of scanner manufacturers and MRI facilities. However, a sufficient sample size is essential as it must contain (1) a representative number of cases and controls from (2) across the lifespan in (3) participants of both sexes, (4) scans at each MRI facility and on scanners from each manufacturer. This is required to avoid possible confounds from interactions of scanner type and age, scanner type and diagnosis, and scanner type and sex. A second option is to generate simulated data from a large enough sample of participants, sites, and MRI facilities. The simulated data could be generated by adding characteristic noise, covariance, and bias profiles for each scanner manufacturer and each MRI facility. The simulated data could then be harmonized with several tools of interest to determine the method that produces data that most closely resembles the pre-noised data. Along the same lines, the post-harmonization data and the pre-noised data could be modeled for case-control effects, age effects, and interaction effects. The results of statistical modeling on post-harmonization datasets could be compared to the results from modeling the pre-noised dataset. The harmonization method that leads to results that most closely resemble the results obtained from modeling the pre-noised data would be deemed most faithful to the ground truth. Scanning an appropriate phantom may add value to ascertaining the ground truth, but is unlikely to add value to characterizing the role of age, sex, and diagnosis on harmonization methods.

While our study focused on 4 widely adopted harmonization methods, these represent only a small number in a large array of available methods. There has been a recent explosion in methods that apply machine learning and other advanced multivariate techniques to tackle harmonization. Machine learning methods, including deep-learning approaches, have been developed in recent years to harmonize neuroimaging data without *a priori* hypotheses about data distributions (Blumberg et al., 2019; Dinsdale et al., 2021; Liu et al., 2021; Moyer et al., 2020; Ning et al., 2020; Tax et al., 2019). A universal machine learning method that is capable of harmonizing data across sites, time, and imaging modalities may enhance our ability to use both existing data and yet to be acquired data (Zuo et al., 2021). For instance, a machine learning harmonization method proposed by Blumberg et al. (2019) applies *multi-task learning*.

From a biological perspective, multi-task learning is inspired by human learning that applies knowledge of performing a known task to new tasks that are related to the known task. Similarly, a subset of harmonization solutions may be found by learning scanner invariant representations while simultaneously maintaining performance on the main task of interest, which include representations of the data that are uninformative about the scanner or the site where imaging data was collected. These representations and the mappings between them may then be manipulated to provide image reconstructions that are minimally informative of their original collection site (Blumberg et al., 2019). This method has several advantages over regression-based methods, including a practical implementation that does not require paired data from a traveling phantom as training input, and extensibility to a multi-site case (Huynh et al., 2019).

Other machine learning developments feature *domain adaptation* techniques (Robinson et al., 2020) that provide immunity against test case data which departs appreciably from the training data, referred to as *domain shift* (Chen et al., 2020a). Domain adaptation techniques attempt to address domain shift by finding a feature space that performs a prescribed main task while being invariant to the domain of the data. Therefore, domain adaptation should be well-suited to MRI data harmonization by creating features that are indiscriminate with respect to the scanner, but correctly discriminate with respect to the data features of interest such as case-control status, age, etc. (Chen et al., 2020b). Convolutional neural networks (CNN), which are popular and well-adapted to vision problems, have also been deployed for data harmonization with demonstrated success at age prediction, although CNN performance can be susceptible to registration related artifacts (Dinsdale et al., 2021). Ning et al. (2020) evaluated 19 algorithms that were developed to harmonize cross-scanner and cross-protocol multi-shell diffusion MRI data acquired from the participant sample on two scanners with different maximum gradient strength using two protocols. The algorithms use various signal representation approaches and computational tools, such as rotational invariant spherical harmonics, deep neural networks, and hybrid biophysical and statistical approaches.

The dawn of the big data age has heralded the need for harmonization methods that operate well beyond neuroimaging data to flexibly and extensibly harmonize manifold data types from social media, mobile devices, and sensors (Agarwal et al., 2013; Davatzikos, 2019). The rapid proliferation of data harmonization methods and the ubiquity of machine learning applications will require careful vetting and rigorous comparisons between competing methods using standard criteria for ascertaining harmonization success. The urgent goal of embracing *open science* will be facilitated by developing advanced harmonization methods (Foster and Deardorff, 2017).

### Limitations

There are two major limitations in the present study. Firstly, we investigated age-related changes in cortical thickness. However, only one scan was administered on each participant in this dataset. New approaches have been developed to harmonize data across scanners and sites as well as longitudinal visits (Beer et al., 2020; Dewey et al., 2019). Age-related cortical thinning estimated by one longitudinal study design was 3 times greater than cortical thinning from a cross-sectional study (Rast et al., 2018). Secondly, we only investigated cortical thickness, which is one of many brain measures that is disturbed in neuropsychiatric disorders. Further studies should fully investigate the performance of harmonization methods on multi- modal neuroimaging data with various anatomical, diffusion, functional, and clinical/behavioral measures.

## Conclusion

ComBat-GAM relative to LME_INT_, LME_INT+SLP_, and ComBat is more sensitive for detecting regions showing significant case-control differences in cortical thickness, and case-control differences in age-related effect on cortical thickness. ComBat-GAM led to larger estimates of cortical thickness reductions, smaller age-related declines, and lower female to male contrast in cases compared to controls. ComBat-GAM also led to greater estimates of age-related declines in cortical thickness in both cases and controls. Our results support using ComBat-GAM to harmonize cortical thickness data across study sites to increase statistical power.

## Acknowledgments

DoD W81XWH-10-1-0925; Center for Brain and Behavior Research Pilot Grant; South Dakota Governor’s Research Center Grant; CX001600 VA CDA; NHMRC Program Grant #1073041; R01 MH111671; VISN6 MIRECC; German Research Foundation grant to J. K. Daniels (DA 1222/4-1 and WA 1539/8-2); VA RR&D 1IK2RX000709; NIMH R01-MH043454; NIMH K01- MH122774; NIMH K01 MH118428-01 (Suarez-Jimenez); 5U01AA021681-08; K24MH71434; K24 DA028773; R01 MH63407; K99NS096116; VA RR&D 1K1RX002325; VA RR&D 1K2RX002922; MH101380; ZonMw, the Netherlands organization for Health Research and Development grant to Miranda Olff (40-00812-98-10041); Academic Medical Center Research Council grant to Miranda Olff (110614); VA CSR&D 1IK2CX001680; VISN17 Center of Excellence pilot funding; NIMH R01MH105535; NIMH 1R21MH102634; German Federal Ministry of Education and Research (BMBF RELEASE 01KR1303A); German Research Society (Deutsche Forschungsgemeinschaft, DFG; SFB/TRR 58: C06, C07); R01MH117601; R01AG059874; MJFF 14848; MH098212; MH071537; M01RR00039; UL1TR000454; HD071982; HD085850; R21MH112956; Anonymous Women’s Health Fund; Kasparian Fund; Trauma Scholars Fund; Barlow Family Fund; NIMH K01 MH118467; Department of Veterans Affairs via support for the National Center for PTSD; NIAAA via its support for (P50) Center for the Translational Neuroscience of Alcohol; NCATS via its support of (CTSA) Yale Center for Clinical Investigation; NIH R01 MH106574; F32MH109274; NIMH 1R21MH102634; R01MH113574; R01-MH103291; BOF 2-4 year project to Sven C. Mueller (01J05415); R01MH105355; Dana Foundation (to Dr. Nitschke); the University of Wisconsin Institute for Clinical and Translational Research; a National Science Foundation Graduate Research Fellowship (to Dr. Grupe); the National Institute of Mental Health (NIMH) R01 MH63407 (to De Bellis), R01 AA12479 (to De Bellis), and R01 MH61744 (to De Bellis); R01- MH043454 and T32-MH018931 (to Dr. Davidson); core grant to the Waisman Center from the National Institute of Child Health and Human Development (P30-HD003352); NIMH K23MH112873; Veterans Affairs Merit Review Program (10/01/08 – 09/30/13); L30 MH114379; South African Medical Research Council “SHARED ROOTS” Flagship Project; Grant MRC-RFA-FSP-01-2013/SHARED ROOTS; South African Research Chair in PTSD from the Department of Science and Technology and the National Research Foundation; US Department of Defense Grant W81XWH08-2-0159 (PI: Stein, Murray B); VA RR&D I01RX000622; CDMRP W81XWH-08–2–0038; South African Medical Research Council; NARSAD Young Investigator; Department of Defense award number W81XWH-12-2-0012; ENIGMA was also supported in part by NIH U54 EB020403 from the Big Data to Knowledge (BD2K) program; R56AG058854; R01MH116147;; P41 EB015922; 1R01MH110483; 1R21 MH098198; R01MH105355-01A. The views expressed in this article are those of the authors and do not necessarily reflect the position or policy of the Department of Veterans Affairs, the United States Government, or any other funding sources listed here.

## Conflicts of Interest

Dr. Abdallah has served as a consultant, speaker and/or on advisory boards for FSV7, Lundbeck, Psilocybin Labs, Genentech and Janssen, and editor of Chronic Stress for Sage Publications, Inc.; he has filed a patent for using mTOR inhibitors to augment the effects of antidepressants (filed on August 20, 2018). Dr. Davidson is the founder and president of, and serves on the board of directors for, the non-profit organization Healthy Minds Innovations, Inc. Dr. Jahanshad, Dr. Thompson and Dr. Ching received partial research support from Biogen, Inc. (Boston, USA) for research unrelated to the content of this manuscript. Dr. Krystal is a consultant for AbbVie, Inc., Amgen, Astellas Pharma Global Development, Inc., AstraZeneca Pharmaceuticals, Biomedisyn Corporation, Bristol-Myers Squibb, Eli Lilly and Company, Euthymics Bioscience, Inc., Neurovance, Inc., FORUM Pharmaceuticals, Janssen Research & Development, Lundbeck Research USA, Novartis Pharma AG, Otsuka America Pharmaceutical, Inc., Sage Therapeutics, Inc., Sunovion Pharmaceuticals, Inc., and Takeda Industries; is on the Scientific Advisory Board for Lohocla Research Corporation, Mnemosyne Pharmaceuticals, Inc., Naurex, Inc., and Pfizer; is a stockholder in Biohaven Pharmaceuticals; holds stock options in Mnemosyne Pharmaceuticals, Inc.; holds patents for Dopamine and Noradrenergic Reuptake Inhibitors in Treatment of Schizophrenia, US Patent No. 5,447,948 (issued September 5, 1995), and Glutamate Modulating Agents in the Treatment of Mental Disorders, U.S. Patent No. 8,778,979 (issued July 15, 2014); and filed a patent for Intranasal Administration of Ketamine to Treat Depression. U.S. Application No. 14/197,767 (filed on March 5, 2014); US application or Patent Cooperation Treaty international application No. 14/306,382 (filed on June 17, 2014); Filed a patent for using mTOR inhibitors to augment the effects of antidepressants (filed on August 20, 2018). Dr. Schmahl is a consultant for Boehringer Ingelheim International GmbH. Dr. Stein has received research grants and/or consultancy honoraria from Lundbeck and Sun. Dr. Lebois reports unpaid membership on the Scientific Committee for the International Society for the Study of Trauma and Dissociation (ISSTD) and spousal license payment for Vanderbilt IP from Acadia Pharmaceuticals unrelated to the topic of this manuscript. All other authors have no conflicts of interest to declare.

## References

Agarwal, P., Shroff, G., Malhotra, P., 2013. Approximate Incremental Big-Data Harmonization. 2013 IEEE International Congress on Big Data, 118–125.

Beer, J.C., Tustison, N.J., Cook, P.A., Davatzikos, C., Sheline, Y.I., Shinohara, R.T., Linn, K.A., Initia, A.D.N., 2020. Longitudinal ComBat: A method for harmonizing longitudinal multi-scanner imaging data. Neuroimage 220.

Blumberg, S.B., Palombo, M., Khoo, C.S., Tax, C.M.W., Tanno, R., Alexander, D.C., 2019. Multi-stage Prediction Networks for Data Harmonization. Medical Image Computing and Computer Assisted Intervention - Miccai 2019, Pt Iv 11767, 411–419.

Boedhoe, P.S., Schmaal, L., Abe, Y., Ameis, S.H., Arnold, P.D., Batistuzzo, M.C., Benedetti, F., Beucke, J.C., Bollettini, I., Bose, A., Brem, S., Calvo, A., Cheng, Y., Cho, K.I., Dallaspezia, S., Denys, D., Fitzgerald, K.D., Fouche, J.P., Gimenez, M., Gruner, P., Hanna, G.L., Hibar, D.P., Hoexter, M.Q., Hu, H., Huyser, C., Ikari, K., Jahanshad, N., Kathmann, N., Kaufmann, C., Koch, K., Kwon, J.S., Lazaro, L., Liu, Y., Lochner, C., Marsh, R., Martinez-Zalacain, I., Mataix-Cols, D., Menchon, J.M., Minuzzi, L., Nakamae, T., Nakao, T., Narayanaswamy, J.C., Piras, F., Piras, F., Pittenger, C., Reddy, Y.C., Sato, J.R., Simpson, H.B., Soreni, N., Soriano-Mas, C., Spalletta, G., Stevens, M.C., Szeszko, P.R., Tolin, D.F., Venkatasubramanian, G., Walitza, S., Wang, Z., van Wingen, G.A., Xu, J., Xu, X., Yun, J.Y., Zhao, Q., Group, E.O.W., Thompson, P.M., Stein, D.J., van den Heuvel, O.A., 2017. Distinct Subcortical Volume Alterations in Pediatric and Adult OCD: A Worldwide Meta- and Mega-Analysis. Am J Psychiatry 174, 60–69.

Brown, S.A., Brumback, T.Y., Tomlinson, K., Cummins, K., Thompson, W.K., Nagel, B.J., De Bellis, M.D., Hooper, S.R., Clark, D.B., Chung, T., Hasler, B.P., Colrain, I.M., Baker, F.C., Prouty, D., Pfefferbaum, A., Sullivan, E.V., Pohl, K.M., Rohlfing, T., Nichols, B.N., Chu, W.W., Tapert, S.F., 2015. The National Consortium on Alcohol and Neuro-Development in Adolescence (NCANDA): A Multisite Study of Adolescent Development and Substance Use. Journal of Studies on Alcohol and Drugs 76, 895–908.

Chen, C., Zheng, Z., Ding, X., Huang, Y., Dou, Q., 2020a. Harmonizing Transferability and Discriminability for Adapting Object Detectors. Proceedings of the IEEE/CVF Conference on Computer Vision and Pattern Recognition, 8869–8878.

Chen, G., Nash, T.A., Reding, K.M., Kohn, P.D., Wei, S.-M., Gregory, M.D., Eisenberg, D.P., Cox, R.W., Berman, K.F., Kippenhan, J.S., 2020b. Beyond linearity in neuroimaging: Capturing nonlinear relationships with application to longitudinal studies. bioRxiv, 2020.2011.2001.363838.

Davatzikos, C., 2019. Machine learning in neuroimaging: Progress and challenges. Neuroimage 197, 652–656.

De Bellis, M.D., Hooper, S.R., Chen, S.D., Provenzale, J.M., Boyd, B.D., Glessner, C.E., MacFall, J.R., Payne, M.E., Rybczynski, R., Woolley, D.P., 2015. Posterior structural brain volumes differ in maltreated youth with and without chronic posttraumatic stress disorder. Development and Psychopathology 27, 1555–1576.

Dennis, E.L., Baron, D., Bartnik-Olson, B., Caeyenberghs, K., Esopenko, C., Hillary, F.G., Kenney, K., Koerte, I.K., Lin, A.P., Mayer, A.R., Mondello, S., Olsen, A., Thompson, P.M., Tate, D.F., Wilde, E.A., 2020. ENIGMA brain injury: Framework, challenges, and opportunities. Human Brain Mapping.

Destrieux, C., Fischl, B., Dale, A., Halgren, E., 2010. Automatic parcellation of human cortical gyri and sulci using standard anatomical nomenclature. Neuroimage 53, 1–15.

Dewey, B.E., Zhao, C., Reinhold, J.C., Carass, A., Fitzgerald, K.C., Sotirchos, E.S., Saidha, S., Oh, J., Pham, D.L., Calabresi, P.A., van Zijl, P.C.M., Prince, J.L., 2019. DeepHarmony: A deep learning approach to contrast harmonization across scanner changes. Magn Reson Imaging 64, 160–170.

Dinsdale, N.K., Jenkinson, M., Namburete, A.I.L., 2021. Deep learning-based unlearning of dataset bias for MRI harmonisation and confound removal. Neuroimage 228, 117689.

Favre, P., Pauling, M., Stout, J., Hozer, F., Sarrazin, S., Abe, C., Alda, M., Alloza, C., Alonso-Lana, S., Andreassen, O.A., Baune, B.T., Benedetti, F., Busatto, G.F., Canales-Rodriguez, E.J., Caseras, X., Chaim-Avancini, T.M., Ching, C.R.K., Dannlowski, U., Deppe, M., Eyler, L.T., Fatjo-Vilas, M., Foley, S.F., Grotegerd, D., Hajek, T., Haukvik, U.K., Howells, F.M., Jahanshad, N., Kugel, H., Lagerberg, T.V., Lawrie, S.M., Linke, J.O., McIntosh, A., Melloni, E.M.T., Mitchell, P.B., Polosan, M., Pomarol-Clotet, E., Repple, J., Roberts, G., Roos, A., Rosa, P.G.P., Salvador, R., Sarro, S., Schofield, P.R., Serpa, M.H., Sim, K., Stein, D.J., Sussmann, J.E., Temmingh, H.S., Thompson, P.M., Verdolini, N., Vieta, E., Wessa, M., Whalley, H.C., Zanetti, M.V., Leboyer, M., Mangin, J.F., Henry, C., Duchesnay, E., Houenou, J., Group, E.B.D.W., 2019. Widespread white matter microstructural abnormalities in bipolar disorder: evidence from mega- and meta-analyses across 3033 individuals. Neuropsychopharmacology 44, 2285–2293.

Fortin, J.P., Cullen, N., Sheline, Y.I., Taylor, W.D., Aselcioglu, I., Cook, P.A., Adams, P., Cooper, C., Fava, M., McGrath, P.J., McInnis, M., Phillips, M.L., Trivedi, M.H., Weissman, M.M., Shinohara, R.T., 2018. Harmonization of cortical thickness measurements across scanners and sites. Neuroimage 167, 104–120.

Fortin, J.P., Parker, D., Tunc, B., Watanabe, T., Elliott, M.A., Ruparel, K., Roalf, D.R., Satterthwaite, T.D., Gur, R.C., Gur, R.E., Schultz, R.T., Verma, R., Shinohara, R.T., 2017. Harmonization of multi-site diffusion tensor imaging data. Neuroimage 161, 149–170.

Foster, E.D., Deardorff, A., 2017. Open Science Framework (OSF). Journal of the Medical Library Association 105, 203.

Frangou, S., Modabbernia, A., Williams, S.C.R., Papachristou, E., Doucet, G.E., Agartz, I., Aghajani, M., Akudjedu, T.N., Albajes-Eizagirre, A., Alnaes, D., Alpert, K.I., Andersson, M., Andreasen, N.C., Andreassen, O.A., Asherson, P., Banaschewski, T., Bargallo, N., Baumeister, S., Baur-Streubel, R., Bertolino, A., Bonvino, A., Boomsma, D.I., Borgwardt, S., Bourque, J., Brandeis, D., Breier, A., Brodaty, H., Brouwer, R.M., Buitelaar, J.K., Busatto, G.F., Buckner, R.L., Calhoun, V., Canales-Rodriguez, E.J., Cannon, D.M., Caseras, X., Castellanos, F.X., Cervenka, S., Chaim-Avancini, T.M., Ching, C.R.K., Chubar, V., Clark, V.P., Conrod, P., Conzelmann, A., Crespo-Facorro, B., Crivello, F., Crone, E.A., Dale, A.M., Dannlowski, U., Davey, C., de Geus, E.J.C., de Haan, L., de Zubicaray, G.I., den Braber, A., Dickie, E.W., Di Giorgio, A., Doan, N.T., Dorum, E.S., Ehrlich, S., Erk, S., Espeseth, T., Fatouros-Bergman, H., Fisher, S.E., Fouche, J.P., Franke, B., Frodl, T., Fuentes-Claramonte, P., Glahn, D.C., Gotlib, I.H., Grabe, H.J., Grimm, O., Groenewold, N.A., Grotegerd, D., Gruber, O., Gruner, P., Gur, R.E., Gur, R.C., Hahn, T., Harrison, B.J., Hartman, C.A., Hatton, S.N., Heinz, A., Heslenfeld, D.J., Hibar, D.P., Hickie, I.B., Ho, B.C., Hoekstra, P.J., Hohmann, S., Holmes, A.J., Hoogman, M., Hosten, N., Howells, F.M., Hulshoff Pol, H.E., Huyser, C., Jahanshad, N., James, A., Jernigan, T.L., Jiang, J., Jonsson, E.G., Joska, J.A., Kahn, R., Kalnin, A., Kanai, R., Klein, M., Klyushnik, T.P., Koenders, L., Koops, S., Kramer, B., Kuntsi, J., Lagopoulos, J., Lazaro, L., Lebedeva, I., Lee, W.H., Lesch, K.P., Lochner, C., Machielsen, M.W.J., Maingault, S., Martin, N.G., Martinez-Zalacain, I., Mataix-Cols, D., Mazoyer, B., McDonald, C., McDonald, B.C., McIntosh, A.M., McMahon, K.L., McPhilemy, G., Meinert, S., Menchon, J.M., Medland, S.E., Meyer-Lindenberg, A., Naaijen, J., Najt, P., Nakao, T., Nordvik, J.E., Nyberg, L., Oosterlaan, J., de la Foz, V.O., Paloyelis, Y., Pauli, P., Pergola, G., Pomarol-Clotet, E., Portella, M.J., Potkin, S.G., Radua, J., Reif, A., Rinker, D.A., Roffman, J.L., Rosa, P.G.P., Sacchet, M.D., Sachdev, P.S., Salvador, R., Sanchez-Juan, P., Sarro, S., Satterthwaite, T.D., Saykin, A.J., Serpa, M.H., Schmaal, L., Schnell, K., Schumann, G., Sim, K., Smoller, J.W., Sommer, I., Soriano-Mas, C., Stein, D.J., Strike, L.T., Swagerman, S.C., Tamnes, C.K., Temmingh, H.S., Thomopoulos, S.I., Tomyshev, A.S., Tordesillas-Gutierrez, D., Trollor, J.N., Turner, J.A., Uhlmann, A., van den Heuvel, O.A., van den Meer, D., van der Wee, N.J.A., van Haren, N.E.M., van ’t Ent, D., van Erp, T.G.M., Veer, I.M., Veltman, D.J., Voineskos, A., Volzke, H., Walter, H., Walton, E., Wang, L., Wang, Y., Wassink, T.H., Weber, B., Wen, W., West, J.D., Westlye, L.T., Whalley, H., Wierenga, L.M., Wittfeld, K., Wolf, D.H., Worker, A., Wright, M.J., Yang, K., Yoncheva, Y., Zanetti, M.V., Ziegler, G.C., Karolinska Schizophrenia, P., Thompson, P.M., Dima, D., 2021. Cortical thickness across the lifespan: Data from 17,075 healthy individuals aged 3-90 years. Human Brain Mapping.

Hatton, S.N., Huynh, K.H., Bonilha, L., Abela, E., Alhusaini, S., Altmann, A., Alvim, M.K.M., Balachandra, A.R., Bartolini, E., Bender, B., Bernasconi, N., Bernasconi, A., Bernhardt, B., Bargallo, N., Caldairou, B., Caligiuri, M.E., Carr, S.J.A., Cavalleri, G.L., Cendes, F., Concha, L., Davoodi-bojd, E., Desmond, P.M., Devinsky, O., Doherty, C.P., Domin, M., Duncan, J.S., Focke, N.K., Foley, S.F., Gambardella, A., Gleichgerrcht, E., Guerrini, R., Hamandi, K., Ishikawa, A., Keller, S.S., Kochunov, P.V., Kotikalapudi, R., Kreilkamp, B.A.K., Kwan, P., Labate, A., Langner, S., Lenge, M., Liu, M., Lui, E., Martin, P., Mascalchi, M., Moreira, J.C.V., Morita-Sherman, M.E., O’Brien, T.J., Pardoe, H.R., Pariente, J.C., Ribeiro, L.F., Richardson, M.P., Rocha, C.S., Rodriguez-Cruces, R., Rosenow, F., Severino, M., Sinclair, B., Soltanian-Zadeh, H., Striano, P., Taylor, P.N., Thomas, R.H., Tortora, D., Velakoulis, D., Vezzani, A., Vivash, L., von Podewils, F., Vos, S.B., Weber, B., Winston, G.P., Yasuda, C.L., Zhu, A.H., Thompson, P.M., Whelan, C.D., Jahanshad, N., Sisodiya, S.M., McDonald, C.R., 2020. White matter abnormalities across different epilepsy syndromes in adults: an ENIGMA-Epilepsy study. Brain 143, 2454–2473.

Hicks, R., Giacino, J., Harrison-Felix, C., Manley, G., Valadka, A., Wilde, E.A., 2013. Progress in Developing Common Data Elements for Traumatic Brain Injury Research: Version Two - The End of the Beginning. Journal of Neurotrauma 30, 1852–1861.

Hofer, E., Roshchupkin, G.V., Adams, H.H.H., Knol, M.J., Lin, H.H., Li, S., Zare, H., Ahmad, S., Armstrong, N.J., Satizabal, C.L., Bernard, M., Bis, J.C., Gillespie, N.A., Luciano, M., Mishra, A., Scholz, M., Teumer, A., Xia, R., Jian, X.Q., Mosley, T.H., Saba, Y., Pirpamer, L., Seiler, S., Becker, J.T., Carmichael, O., Rotter, J.I., Psaty, B.M., Lopez, O.L., Amin, N., van der Lee, S.J., Yang, Q., Himali, J.J., Maillard, P., Beiser, A.S., DeCarli, C., Karama, S., Lewis, L., Harris, M., Bastin, M.E., Deary, I.J., Veronica Witte, A., Beyer, F., Loeffler, M., Mather, K.A., Schofield, P.R., Thalamuthu, A., Kwok, J.B., Wright, M.J., Ames, D., Trollor, J., Jiang, J.Y., Brodaty, H., Wen, W., Vernooij, M.W., Hofman, A., Uitterlinden, A.G., Niessen, W.J., Wittfeld, K., Bulow, R., Volker, U., Pausova, Z., Bruce Pike, G., Maingault, S., Crivello, F., Tzourio, C., Amouyel, P., Mazoyer, B., Neale, M.C., Franz, C.E., Lyons, M.J., Panizzon, M.S., Andreassen, O.A., Dale, A.M., Logue, M., Grasby, K.L., Jahanshad, N., Painter, J.N., Colodro-Conde, L., Bralten, J., Hibar, D.P., Lind, P.A., Pizzagalli, F., Stein, J.L., Thompson, P.M., Medland, S.E., Sachdev, P.S., Kremen, W.S., Wardlaw, J.M., Villringer, A., van Duijn, C.M., Grabe, H.J., Longstreth, W.T., Fornage, M., Paus, T., Debette, S., Ikram, M.A., Schmidt, H., Schmidt, R., Seshadri, S., Consortium, E., 2020. Genetic correlations and genome-wide associations of cortical structure in general population samples of 22,824 adults. Nature Communications 11.

Huynh, K.M., Chen, G., Wu, Y., Shen, D., Yap, P.T., 2019. Multi-Site Harmonization of Diffusion MRI Data via Method of Moments. IEEE Trans Med Imaging 38, 1599–1609.

Johnson, W.E., Li, C., Rabinovic, A., 2007. Adjusting batch effects in microarray expression data using empirical Bayes methods. Biostatistics 8, 118–127.

Koshiyama, D., Miura, K., Nemoto, K., Okada, N., Matsumoto, J., Fukunaga, M., Hashimoto, R., 2020. Neuroimaging studies within Cognitive Genetics Collaborative Research Organization aiming to replicate and extend works of ENIGMA. Human Brain Mapping.

Liu, M., Maiti, P., Thomopoulos, S., Zhu, A., Chai, Y., Kim, H., Jahanshad, N., 2021. Style Transfer Using Generative Adversarial Networks for Multi-Site MRI Harmonization. bioRxiv, 2021.2003.2017.435892.

Logue, M.W., van Rooij, S.J.H., Dennis, E.L., Davis, S.L., Hayes, J.P., Stevens, J.S., Densmore, M., Haswell, C.C., Ipser, J., Koch, S.B.J., Korgaonkar, M., Lebois, L.A.M., Peverill, M., Baker, J.T., Boedhoe, P.S.W., Frijling, J.L., Gruber, S.A., Harpaz-Rotem, I., Jahanshad, N., Koopowitz, S., Levy, I., Nawijn, L., O’Connor, L., Olff, M., Salat, D.H., Sheridan, M.A., Spielberg, J.M., van Zuiden, M., Winternitz, S.R., Wolff, J.D., Wolf, E.J., Wang, X., Wrocklage, K., Abdallah, C.G., Bryant, R.A., Geuze, E., Jovanovic, T., Kaufman, M.L., King, A.P., Krystal, J.H., Lagopoulos, J., Bennett, M., Lanius, R., Liberzon, I., McGlinchey, R.E., McLaughlin, K.A., Milberg, W.P., Miller, M.W., Ressler, K.J., Veltman, D.J., Stein, D.J., Thomaes, K., Thompson, P.M., Morey, R.A., 2018. Smaller Hippocampal Volume in Posttraumatic Stress Disorder: A Multisite ENIGMA-PGC Study: Subcortical Volumetry Results From Posttraumatic Stress Disorder Consortia. Biological Psychiatry 83, 244–253.

Morey, R.A., Haswell, C.C., Hooper, S.R., De Bellis, M.D., 2016. Amygdala, Hippocampus, and Ventral Medial Prefrontal Cortex Volumes Differ in Maltreated Youth with and without Chronic Posttraumatic Stress Disorder. Neuropsychopharmacology 41, 791–801.

Moyer, D., Steeg, G.V., Tax, C.M.W., Thompson, P.M., 2020. Scanner invariant representations for diffusion MRI harmonization. Magnetic Resonance in Medicine 84, 2174–2189.

Mutlu, A.K., Schneider, M., Debbane, M., Badoud, D., Eliez, S., Schaer, M., 2013. Sex differences in thickness, and folding developments throughout the cortex. Neuroimage 82, 200–207.

Ning, L.P., Bonet-Carne, E., Grussu, F., Sepehrband, F., Kaden, E., Veraart, J., Blumberg, S.B., Khoo, C.S., Palombo, M., Kokkinos, I., Alexander, D.C., Coll-Font, J., Scherrer, B., Warfield, S.K., Karayumak, S.C., Rathi, Y., Koppers, S., Weninger, L., Ebert, J., Merhof, D., Moyer, D., Pietsch, M., Christiaens, D., Teixeira, R.A.G., Tournier, J.D., Schilling, K.G., Huo, Y.K., Nath, V., Hansen, C., Blaber, J., Landman, B.A., Zhylka, A., Pluim, J.P.W., Parker, G., Rudrapatna, U., Evans, J., Charron, C., Jones, D.K., Tax, C.M.W., 2020. Cross-scanner and cross-protocol multi-shell diffusion MRI data harmonization: Algorithms and results. Neuroimage 221.

Noble, S., Scheinost, D., Finn, E.S., Shen, X.L., Papademetris, X., McEwen, S.C., Bearden, C.E., Addington, J., Goodyear, B., Cadenhead, K.S., Mirzakhanian, H., Cornblatt, B.A., Olvet, D.M., Mathalon, D.H., McGlashan, T.H., Perkins, D.O., Belger, A., Seidman, L.J., Thermenos, H., Tsuang, M.T., van Erp, T.G.M., Walker, E.F., Hamann, S., Woods, S.W., Cannon, T.D., Constable, R.T., 2017. Multisite reliability of MR-based functional connectivity. Neuroimage 146, 959–970.

Pomponio, R., Erus, G., Habes, M., Doshi, J., Srinivasan, D., Mamourian, E., Bashyam, V., Nasrallah, I.M., Satterthwaite, T.D., Fan, Y., Launer, L.J., Masters, C.L., Maruff, P., Zhuo, C.J., Volzke, H., Johnson, S.C., Fripp, J., Koutsouleris, N., Wolf, D.H., Gur, R., Gur, R., Morris, J., Albert, M.S., Grabe, H.J., Resnick, S.M., Bryan, R.N., Wolk, D.A., Shinohara, R.T., Shou, H.C., Davatzikos, C., 2020. Harmonization of large MRI datasets for the analysis of brain imaging patterns throughout the lifespan. Neuroimage 208.

Radua, J., Vieta, E., Shinohara, R., Kochunov, P., Quide, Y., Green, M.J., Weickert, C.S., Weickert, T., Bruggemann, J., Kircher, T., Nenadic, I., Cairns, M.J., Seal, M., Schall, U., Henskens, F., Fullerton, J.M., Mowry, B., Pantelis, C., Lenroot, R., Cropley, V., Loughland, C., Scott, R., Wolf, D., Satterthwaite, T.D., Tan, Y., Sim, K., Piras, F., Spalletta, G., Banaj, N., Pomarol-Clotet, E., Solanes, A., Albajes-Eizagirre, A., Canales-Rodriguez, E.J., Sarro, S., Di Giorgio, A., Bertolino, A., Stablein, M., Oertel, V., Knochel, C., Borgwardt, S., du Plessis, S., Yun, J.Y., Kwon, J.S., Dannlowski, U., Hahn, T., Grotegerd, D., Alloza, C., Arango, C., Janssen, J., Diaz-Caneja, C., Jiang, W., Calhoun, V., Ehrlich, S., Yang, K., Cascella, N.G., Takayanagi, Y., Sawa, A., Tomyshev, A., Lebedeva, I., Kaleda, V., Kirschner, M., Hoschl, C., Tomecek, D., Skoch, A., van Amelsvoort, T., Bakker, G., James, A., Preda, A., Weideman, A., Stein, D.J., Howells, F., Uhlmann, A., Temmingh, H., Lopez-Jaramillo, C., Diaz-Zuluaga, A., Fortea, L., Martinez-Heras, E., Solana, E., Llufriu, S., Jahanshad, N., Thompson, P., Turner, J., van Erp, T., collaborators, E.C., 2020. Increased power by harmonizing structural MRI site differences with the ComBat batch adjustment method in ENIGMA. Neuroimage 218, 116956.

Rast, P., Kennedy, K.M., Rodrigue, K.M., Robinson, P., Gross, A.L., McLaren, D.G., Grabowski, T., Schaie, K.W., Willis, S.L., 2018. APOEepsilon4 Genotype and Hypertension Modify 8-year Cortical Thinning: Five Occasion Evidence from the Seattle Longitudinal Study. Cerebral Cortex 28, 1934–1945.

Ritchie, S.J., Cox, S.R., Shen, X., Lombardo, M.V., Reus, L.M., Alloza, C., Harris, M.A., Alderson, H.L., Hunter, S., Neilson, E., Liewald, D.C.M., Auyeung, B., Whalley, H.C., Lawrie, S.M., Gale, C.R., Bastin, M.E., McIntosh, A.M., Deary, I.J., 2018. Sex Differences in the Adult Human Brain: Evidence from 5216 UK Biobank Participants. Cerebral Cortex 28, 2959–2975.

Robinson, R., Dou, Q., Castro, D.C., Kamnitsas, K., Groot, M.d., Summers, R.M., Rueckert, D., Glocker, B., 2020. Image-level Harmonization of Multi-Site Data using Image-and-Spatial Transformer Networks. Medical Image Computing and Computer-Assisted Intervention, 710–719.

Santhanam, P., Wilson, S.H., Mulatya, C., Oakes, T.R., Weaver, L.K., 2019. Age-Accelerated Reduction in Cortical Surface Area in United States Service Members and Veterans with Mild Traumatic Brain Injury and Post-Traumatic Stress Disorder. Journal of Neurotrauma 36, 2922–2929.

Savalia, N.K., Agres, P.F., Chan, M.Y., Feczko, E.J., Kennedy, K.M., Wig, G.S., 2017. Motion-related artifacts in structural brain images revealed with independent estimates of in-scanner head motion. Human Brain Mapping 38, 472–492.

Savjani, R.R., Taylor, B.A., Acion, L., Wilde, E.A., Jorge, R.E., 2017. Accelerated Changes in Cortical Thickness Measurements with Age in Military Service Members with Traumatic Brain Injury. Journal of Neurotrauma 34, 3107–3116.

Shalev, A., Liberzon, I., Marmar, C., 2017. Post-Traumatic Stress Disorder. New England Journal of Medicine 376, 2459–2469.

Tax, C.M.W., Grussu, F., Kaden, E., Ning, L.P., Rudrapatna, U., Evans, C.J., St-Jean, S., Leemans, A., Koppers, S., Merhof, D., Ghosh, A., Tanno, R., Alexander, D.C., Zappala, S., Charron, C., Kusmia, S., Linden, D.E.J., Jones, D.K., Veraart, J., 2019. Cross-scanner and cross-protocol diffusion MRI data harmonisation: A benchmark database and evaluation of algorithms. Neuroimage 195, 285–299.

Thompson, P.M., Jahanshad, N., Ching, C.R.K., Salminen, L.E., Thomopoulos, S.I., Bright, J., Baune, B.T., Bertolin, S., Bralten, J., Bruin, W.B., Bulow, R., Chen, J., Chye, Y., Dannlowski, U., de Kovel, C.G.F., Donohoe, G., Eyler, L.T., Faraone, S.V., Favre, P., Filippi, C.A., Frodl, T., Garijo, D., Gil, Y., Grabe, H.J., Grasby, K.L., Hajek, T., Han, L.K.M., Hatton, S.N., Hilbert, K., Ho, T.F.C., Holleran, L., Homuth, G., Hosten, N., Houenou, J., Ivanov, I., Jia, T.Y., Kelly, S., Klein, M., Kwon, J.S., Laansma, M.A., Leerssen, J., Lueken, U., Nunes, A., Neill, J.O., Opel, N., Piras, F., Piras, F., Postema, M.C., Pozzi, E., Shatokhina, N., Soriano-Mas, C., Spalletta, G., Sun, D.Q., Teumer, A., Tilot, A.K., Tozzi, L., van der Merwe, C., Van Someren, E.J.W., van Wingen, G.A., Volzke, H., Walton, E., Wang, L., Winkler, A.M., Wittfeld, K., Wright, M.J., Yun, J.Y., Zhang, G.H., Zhang-James, Y., Adhikari, B.M., Agartz, I., Aghajani, M., Aleman, A., Althoff, R.R., Altmann, A., Andreassen, O.A., Baron, D.A., Bartnik-Olson, B.L., Bas-Hoogendam, J.M., Baskin-Sommers, A.R., Bearden, C.E., Berner, L.A., Boedhoe, P.S.W., Brouwer, R.M., Buitelaar, J.K., Caeyenberghs, K., Cecil, C.A.M., Cohen, R.A., Cole, J.H., Conrod, P.J., De Brito, S.A., de Zwarte, S.M.C., Dennis, E.L., Desrivieres, S., Dima, D., Ehrlich, S., Esopenko, C., Fairchild, G., Fisher, S.E., Fouche, J.P., Francks, C., Frangou, S., Franke, B., Garavan, H.P., Glahn, D.C., Groenewold, N.A., Gurholt, T.P., Gutman, B.A., Hahn, T., Harding, I.H., Hernaus, D., Hibar, D.P., Hillary, F.G., Hoogman, M., Pol, H.H.E., Jalbrzikowski, M., Karkashadze, G.A., Klapwijk, E.T., Knickmeyer, R.C., Kochunov, P., Koerte, I.K., Kong, X.Z., Liew, S.L., Lin, A.L.P., Logue, M.W., Luders, E., Macciardi, F., Mackey, S., Mayer, A.R., McDonald, C.R., McMahon, A.B., Medland, S.E., Modinos, G., Morey, R.A., Mueller, S.C., Mukherjee, P., Namazova-Baranova, L., Nir, T.M., Olsen, A., Paschou, P., Pine, D.S., Pizzagalli, F., Renteria, M.E., Rohrer, J.D., Samann, P.G., Schmaal, L., Schumann, G., Shiroishi, M.S., Sisodiya, S.M., Smit, D.J.A., Sonderby, I.E., Stein, D.J., Stein, J.L., Tahmasian, M., Tate, D.F., Turner, J.A., van den Heuvel, O.A., van der Wee, N.J.A., van der Werf, Y.D., van Erp, T.G.M., van Haren, N.E.M., van Rooij, D., van Velzen, L.S., Veer, I.M., Veltman, D.J., Villalon-Reina, J.E., Walter, H., Whelan, C.D., Wilde, E.A., Zarei, M., Zelman, V., Consortium, E., 2020. ENIGMA and global neuroscience: A decade of large-scale studies of the brain in health and disease across more than 40 countries. Translational Psychiatry 10.

van Rooij, D., Anagnostou, E., Arango, C., Auzias, G., Behrmann, M., Busatto, G.F., Calderoni, S., Daly, E., Deruelle, C., Di Martino, A., Dinstein, I., Duran, F.L.S., Durston, S., Ecker, C., Fair, D., Fedor, J., Fitzgerald, J., Freitag, C.M., Gallagher, L., Gori, I., Haar, S., Hoekstra, L., Jahanshad, N., Jalbrzikowski, M., Janssen, J., Lerch, J., Luna, B., Martinho, M.M., McGrath, J., Muratori, F., Murphy, C.M., Murphy, D.G.M., O’Hearn, K., Oranje, B., Parellada, M., Retico, A., Rosa, P., Rubia, K., Shook, D., Taylor, M., Thompson, P.M., Tosetti, M., Wallace, G.L., Zhou, F., Buitelaar, J.K., 2018. Cortical and Subcortical Brain Morphometry Differences Between Patients With Autism Spectrum Disorder and Healthy Individuals Across the Lifespan: Results From the ENIGMA ASD Working Group. Am J Psychiatry 175, 359–369.

Volkow, N.D., Koob, G.F., Croyle, R.T., Bianchi, D.W., Gordon, J.A., Koroshetz, W.J., Perez-Stable, E.J., Riley, W.T., Bloch, M.H., Conway, K., Deeds, B.G., Dowling, G.J., Grant, S., Howlett, K.D., Matochik, J.A., Morgan, G.D., Murray, M.M., Noronha, A., Spong, C.Y., Wargo, E.M., Warren, K.R., Weiss, S.R.B., 2018. The conception of the ABCD study: From substance use to a broad NIH collaboration. Developmental Cognitive Neuroscience 32, 4–7.

Walhovd, K.B., Fjell, A.M., Giedd, J., Dale, A.M., Brown, T.T., 2017. Through Thick and Thin: a Need to Reconcile Contradictory Results on Trajectories in Human Cortical Development. Cerebral Cortex 27, 1472–1481.

Wang, X., Xie, H., Chen, T., Cotton, A.S., Salminen, L.E., Logue, M.W., Clarke-Rubright, E.K., Wall, J., Dennis, E.L., O’Leary, B.M., Abdallah, C.G., Andrew, E., Baugh, L.A., Bomyea, J., Bruce, S.E., Bryant, R., Choi, K., Daniels, J.K., Davenport, N.D., Davidson, R.J., DeBellis, M., deRoon-Cassini, T., Disner, S.G., Fani, N., Fercho, K.A., Fitzgerald, J., Forster, G.L., Frijling, J.L., Geuze, E., Gomaa, H., Gordon, E.M., Grupe, D., Harpaz-Rotem, I., Haswell, C.C., Herzog, J.I., Hofmann, D., Hollifield, M., Hosseini, B., Hudson, A.R., Ipser, J., Jahanshad, N., Jovanovic, T., Kaufman, M.L., King, A.P., Koch, S.B.J., Koerte, I.K., Korgaonkar, M.S., Krystal, J.H., Larson, C., Lebois, L.A.M., Levy, I., Li, G., Magnotta, V.A., Manthey, A., May, G., McLaughlin, K.A., Mueller, S.C., Nawijn, L., Nelson, S.M., Neria, Y., Nitschke, J.B., Olff, M., Olson, E.A., Peverill, M., Luan Phan, K., Rashid, F.M., Ressler, K., Rosso, I.M., Sambrook, K., Schmahl, C., Shenton, M.E., Sierk, A., Simons, J.S., Simons, R.M., Sponheim, S.R., Stein, M.B., Stein, D.J., Stevens, J.S., Straube, T., Suarez-Jimenez, B., Tamburrino, M., Thomopoulos, S.I., van der Wee, N.J.A., van der Werff, S.J.A., van Erp, T.G.M., van Rooij, S.J.H., van Zuiden, M., Varkevisser, T., Veltman, D.J., Vermeiren, R., Walter, H., Wang, L., Zhu, Y., Zhu, X., Thompson, P.M., Morey, R.A., Liberzon, I., 2020. Cortical volume abnormalities in posttraumatic stress disorder: an ENIGMA-psychiatric genomics consortium PTSD workgroup mega-analysis. Mol Psychiatry.

Yu, M.C., Linn, K.A., Cook, P.A., Phillips, M.L., McInnis, M., Fava, M., Trivedi, M.H., Weissman, M.M., Shinohara, R.T., Sheline, Y.I., 2018. Statistical harmonization corrects site effects in functional connectivity measurements from multi-site fMRI data. Human Brain Mapping 39, 4213–4227.

Zugman, A., Harrewijn, A., Cardinale, E.M., Zwiebel, H., Freitag, G.F., Werwath, K.E., Bas-Hoogendam, J.M., Groenewold, N.A., Aghajani, M., Hilbert, K., Cardoner, N., Porta-Casteras, D., Gosnell, S., Salas, R., Blair, K.S., Blair, J.R., Hammoud, M.Z., Milad, M., Burkhouse, K., Phan, K.L., Schroeder, H.K., Strawn, J.R., Beesdo-Baum, K., Thomopoulos, S.I., Grabe, H.J., Van der Auwera, S., Wittfeld, K., Nielsen, J.A., Buckner, R., Smoller, J.W., Mwangi, B., Soares, J.C., Wu, M.J., Zunta-Soares, G.B., Jackowski, A.P., Pan, P.M., Salum, G.A., Assaf, M., Diefenbach, G.J., Brambilla, P., Maggioni, E., Hofmann, D., Straube, T., Andreescu, C., Berta, R., Tamburo, E., Price, R., Manfro, G.G., Critchley, H.D., Makovac, E., Mancini, M., Meeten, F., Ottaviani, C., Agosta, F., Canu, E., Cividini, C., Filippi, M., Kostic, M., Munjiza, A., Filippi, C.A., Leibenluft, E., Alberton, B.A.V., Balderston, N.L., Ernst, M., Grillon, C., Mujica-Parodi, L.R., van Nieuwenhuizen, H., Fonzo, G.A., Paulus, M.P., Stein, M.B., Gur, R.E., Gur, R.C., Kaczkurkin, A.N., Larsen, B., Satterthwaite, T.D., Harper, J., Myers, M., Perino, M.T., Yu, Q., Sylvester, C.M., Veltman, D.J., Lueken, U., Van der Wee, N.J.A., Stein, D.J., Jahanshad, N., Thompson, P.M., Pine, D.S., Winkler, A.M., 2020. Mega-analysis methods in ENIGMA: The experience of the generalized anxiety disorder working group. Human Brain Mapping.

Zuo, L., Dewey, B.E., Carass, A., Liu, Y., He, Y., Calabresi, P.A., Prince, J.L., 2021. Information-Based Disentangled Representation Learning for Unsupervised MR Harmonization. Springer International Publishing, Cham, pp. 346–359.

