## Supplementary Materials for "A Comparison of Methods to Harmonize Cortical Thickness Measurements Across Scanners and Sites"

**Methods**

*Participants*

Most sites used clinician-administered measures such as the Structured Clinical Interview (SCID) (First, 2015) or Clinician-Administered PTSD Scale (CAPS) (Weathers et al., 2018; Weathers et al., 2001) to ascertain PTSD diagnosis. One site used a psychiatrist diagnosis for PTSD, and a few additional sites used self-report scales such as the PTSD Checklist (PCL) (Weathers et al., 2013). While the majority used DSM-IV criteria, a small subset of sites used DSM-5 criteria.

**Results**

Significant Age x Diagnosis Interaction Detected by ComBat-GAM

Age-related declines in cortical thickness were slower in cases than controls for 5 regions within the DMN, which include the left middle-posterior part of the cingulate gyrus and sulcus, the right marginal branch of the cingulate sulcus, the right superior frontal sulcus in ECN, right inferior temporal areas that include the right medial occipital-temporal sulcus and lingual sulcus, and the right fusiform gyrus.

The linear fits of the age-related distributions of cortical thickness in the 5 regions were shown **Fig. S1** as shown below.

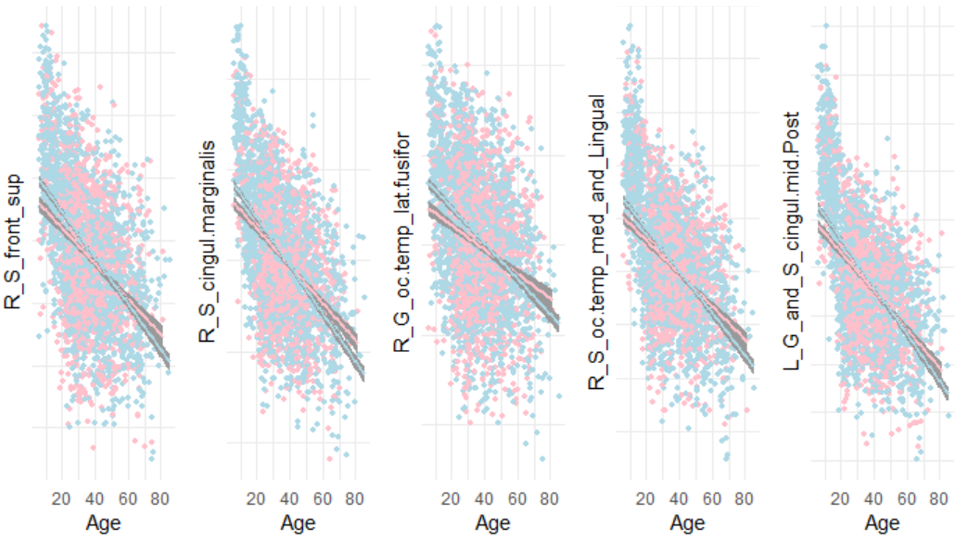

The non-linear fits of the age-related distributions of cortical thickness in the 5 regions were shown **Fig. S2** as shown below.

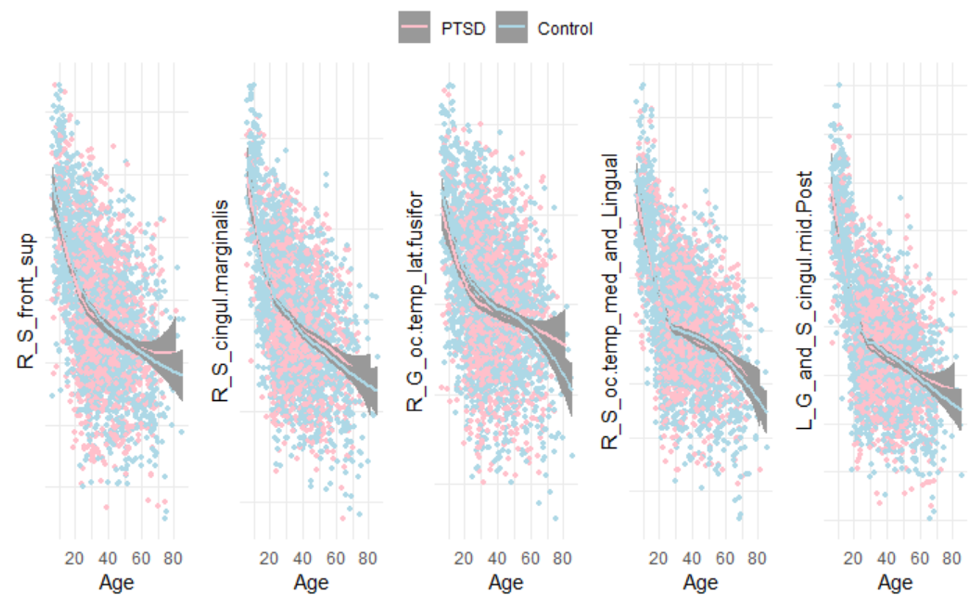

**References**

First, M.B., 2015. Structured Clinical Interview for the DSM (SCID). The Encyclopedia of Clinical Psychology, pp. 1-6.

Weathers, F.W., Blake, D.D., Schnurr, P.P., Kaloupek, D.G., Marx, B.P., Keane, T.M., 2013. The PTSD Checklist for DSM-5 (PCL-5).

Weathers, F.W., Bovin, M.J., Lee, D.J., Sloan, D.M., Schnurr, P.P., Kaloupek, D.G., Keane, T.M., Marx, B.P., 2018. The Clinician-Administered PTSD Scale for DSM-5 (CAPS-5): Development and Initial Psychometric Evaluation in Military Veterans. Psychological Assessment 30, 383-395.

Weathers, F.W., Keane, T.M., Davidson, J.R.T., 2001. Clinician-administered PTSD scale: A review of the first ten years of research. Depression and Anxiety 13, 132-156.
